# Biosensor Cell Array Reveals Temporal GABA Secretion Dynamics from Pancreatic Islets

**DOI:** 10.64898/2026.03.31.715660

**Authors:** Austin E. Stis, Charles S. Lazimi, Sandra M. Ferreira, Alexandra E. Cuaycal, Dylan Smurlick, D. Walker Hagan, Taylor Nakayama, Sunil P. Gandhi, Emery Smith, Timothy P. Spicer, Edward A. Phelps

## Abstract

Pancreatic beta cells have the unique function of synthesizing and secreting high amounts of the inhibitory neurotransmitter γ-aminobutyric acid (GABA). The mechanism of GABA secretion, whether vesicular or channel-mediated, is debated. Our study reveals surprising temporal complexity in the pattern of islet GABA secretion. We used insulin secretion modulators to demonstrate that GABA release is not directly correlated with insulin secretion. VGAT reporter mice also showed that beta cells do not express the requisite vesicular GABA transporter (VGAT) for vesicular GABA release. Instead, GABA is secreted from the cytosol in pulses by the LRRC8A/D isoform of the volume regulatory anion channel (VRAC). We further demonstrate the dynamic coordination of GABA release with calcium influx in beta cells and dependence on beta cell depolarization. These results suggest a model where GABA is released during the peaks of beta cell calcium oscillations to provide feedback which strengthens and reinforces the oscillation waveform.

## Introduction

Pancreatic beta cells are the only non-neural cell type that synthesizes and secretes large quantities of the inhibitory neurotransmitter γ-aminobutyric acid (GABA)^1^. GABA is synthesized by the enzyme glutamate decarboxylase (GAD) which exists in two isoforms: 67 kDa isoform GAD67 (*GAD1*) and 65 kDa isoform GAD65 (*GAD2*). There are species-specific differences in the expression of GAD isoform in beta cells. Human beta cells only express GAD65, mouse beta cells express mostly GAD67 with trace amounts of GAD65, and rat beta cells express both isoforms of GAD^2, 3^. Autoantibodies specific to GAD65 are one of the four major diagnostic autoantibodies in type 1 diabetes^4, 5^.

There is a long history of evidence that GABA is both an autocrine and paracrine signaling molecule in the islet. In beta cells, GABA has been described as alternatively excitatory or inhibitory depending on the experimental concentrations. In low glucose conditions up to 6 mM, GABA has been shown by some studies to be excitatory to beta cells, causing an increase in insulin secretion by helping depolarize the plasma membrane^6, 7, 8^. However, at high glucose concentrations where the beta cell is already excited, GABA is inhibitory and decreases insulin secretion by hyperpolarizing the plasma membrane^7^. Other studies have shown that GABA is always inhibitory^9, 10, 11, 12^. The action of GABA on beta cells depends on the function of the GABA_A_R, a ligand-gated Cl^-^ channel and GABA_B_R, a metabotropic G_i/o_-coupled receptor^13, 14, 15, 16, 17^. Some studies indicate that human beta cells do not form functional GABA_B_R complexes under basal conditions^14^. Immune cells are also negatively regulated by GABA^18, 19, 20^. GABA content becomes depleted in remaining beta cells in islets from donors with type 1 diabetes, which has been hypothesized to leave these islets more vulnerable to autoimmunity^9^.

Multiple pathways have been described for GABA secretion from pancreatic beta cells. In neurons, GABA is synthesized in the cytoplasm of the presynaptic neuron by both GAD65 and GAD67 from glutamate^21^. The vesicular GABA transporter (VGAT), also known as the vesicular inhibitory amino acid transporter (VIAAT), gene name *SLC32A1*, is located on synaptic vesicles at the synapses of the presynaptic neuron and is responsible for the accumulation of GABA and glycine into the lumen of synaptic vesicles^21, 22, 23^. VGAT transports GABA into the synaptic vesicles, which upon depolarization of the presynaptic neuron release GABA into the synaptic cleft^21^.

Early work on the beta cell GABA system concluded that GABA is secreted from synaptic-like microvesicles (analogous to synaptic vesicles in neurons) or co-secreted with insulin from large dense-core vesicles^6, 24^. Braun et al., showed that islet GABA secretion is triggered by glucose or patch-pipetted infusion of free Ca^2+^, suggesting that GABA is co-released with insulin^6, 24^. However, Braun et al., also described a high background secretion of GABA directly from the cytosol that was non-vesicular and independent of glucose^6, 9^. In contrast, our prior work showed that the majority of islet GABA secretion occurs directly from the cytosol, is unregulated by glucose, and is released in periodic pulses^25^. Thus, the understanding of the mechanism, physiological triggers for islet GABA secretion, and its coordination with beta cell excitation requires further study to unify these different findings.

A vesicular release mechanism for GABA in beta cells, either from insulin granules or synaptic-like microvesicles, would likely require VGAT expression. The data for VGAT expression in islets is inconsistent. VGAT has been previously identified in rat beta cells^26, 27^ and a rat beta cell line^28^ by western blots of islet lysate and immunostaining of pancreas tissue sections. Other studies and our own observations show immunoreactivity for VGAT only in the rat islet mantle alpha cells and no staining in rat beta cells^9, 29^. In human beta cells, one study showed positive VGAT immunostaining in human pancreas^28^, while we found the majority of human beta cells did not stain positively for VGAT^9^. There are no other known high-affinity vesicular GABA transporters that could substitute for VGAT. The vesicular monoamine transporter 2 (VMAT2) is a low-affinity vesicular GABA transporter^30, 31, 32, 33^ and is expressed in pancreatic beta cells^34,35^. However, VMAT2 does not transport GABA under physiological conditions^33^. Thus, an explanation for observations of vesicular release of GABA in pancreatic beta cells is unresolved and requires additional confirmation.

We previously sought to determine if membrane channels contribute to GABA efflux directly from the cytosol in beta cells^9^. We looked for expression of known GABA permeant channels, including the volume regulatory anion channels (VRAC)^36^. VRAC is an osmo-sensitive Cl^-^ channel that controls cell volume. Multiple VRAC isoforms are formed as hetero-hexamer complexes of six subunits of LRRC8 family proteins: LRRC8A, B, C, D, or E. The subunit LRRC8A (also known as SWELL1) is obligatory for functional VRAC while its heteromeric partners LRRC8B-E confer functional heterogeneity^36, 37^. In addition, one VRAC isoform (LRRC8A/D) is specialized for permeability to cytosolic organic osmolytes such as GABA and taurine^36, 38, 39, 40^, while the other isoforms are more specific for Cl^-^. Confirmation of the expression of LRRC8A, B, and D in primary beta cells by rt-PCR and western blot has been published by us and others^9, 41, 42, 43, 44^. We showed that LRRC8A is required for non-vesicular release of GABA from pancreatic beta cells and that this secretion has a regular pulsatile pattern. Kang et al., showed that beta cells produce LRRC8A-dependent channel currents in response to hypotonic stimulus or glucose^42^, although we previously did not observe a glucose-dependence of GABA release^25^. Thus, glucose dependence of GABA secretion from the specific LRRC8A/D VRAC isoform requires additional study.

Here, we have introduced multiple unresolved questions surrounding the islet GABA system: (1) The evidence for the expression of VGAT in beta cells is ambiguous; (2) The effect of glucose or cytosolic calcium ion concentration ([Ca^2+^]_i_) on GABA release is inconsistently reported; and (3) The physiological trigger for non-vesicular islet GABA secretion has not been revealed. These results presented in this study are intended to better unify the understanding of the islet GABA system. We confirmed that VGAT is not detected in beta cells using a VGAT reporter mouse, GABA secretion triggered by glucose depends on the timescale of the observation, non-vesicular GABA secretion occurs via the LRRC8A/D isoform of VRAC, and pulsatile GABA release events are closely coordinated with islet [Ca^2+^]_i_ oscillations.

## Methods

### Gad ^βKO^ Mouse Generation

*Gad1^loxp^::Gad2^loxp^::Rosa26^tdTomato^*mice (B6.Cg-*Gad2^tm1Rpa^ Gad1^tm1Rpa^ Gt(ROSA)26Sor^tm14(CAG-tdTomato)Hze^*/RpaJ) (JAX # 031800) were generated in Richard Palmiter’s lab at the University of Washington^45^ and provided by Dr. Qi Wu’s lab at Baylor University College of Medicine were crossed with mice expressing Cre recombinase in beta cells (*Ins1^cre^,* B6(Cg)-*Ins1^tm1.1(cre)Thor^*/J)^46^ (JAX # 026801) to generate the *Gad* ^βKO^ mice. All offspring were genotyped post-wean using tail snips and custom primers generated by Transnetyx specific to each *Gad1^loxp^, Gad2^loxp^, Rosa26^tdTomato^,* and *Ins1^cre^* sequence. Ins-cre mice were used as controls. All experiments performed here were previously approved by the Institutional Animal Care and Use Committee at the University of Florida (202400000147).

### Mouse Islet Isolation

All experimental protocols using transgenic mice were approved by the University of Florida Animal Care and Use Committee. Experimental protocols using transgenic mouse islets were approved by the University of Florida Animal Care and Use Committee. Mice were euthanized by deep isoflurane anesthesia followed by cervical dislocation according to approved protocols. The pancreases of C57Bl/6J, Ins-cre, or *Gad* ^βKO^ mice were perfused via the common bile duct with 2–3 mL HBSS containing collagenase type V (0.6 mg mL^−1^, Sigma C9263), removed into a 15 mL conical tube, and digested at 37 °C for 17 min. The enzyme digestion was quenched with 10 mL of cold HBSS (Gibco 14025092), 10% FBS (Cytiva SH30071), 1% pen/strep (Gibco 15140-122), and 2% 1 M Hepes buffer (Fisher BP299100). Digested pancreas was shaken and then spun down at 200 x g for 2 minutes at 4 °C. Supernatant was discarded, and the pellet was resuspended in 10 mL of cold HBSS with 10% FBS, 1% pen/strep, and 2% 1 M HEPES. Islets were hand-picked using a stereomicroscope (Leica M80). Islets were cultured overnight in RPMI 1640 (Gibco 11875-093) with 10% FBS, and 1% pen/strep at 37 °C.

### Human Donor Islets

Use of cadaveric human pancreatic islets has been approved and classified as “Non-Human Subjects” by the University of Florida Institutional Review Board (IRB #201702860). Isolated and purified human islets were obtained from 3 non-diabetic donors from NIDDK-funded Integrated Islet Distribution Program (IIDP) at City of Hope for GABA secretion static incubations. Isolated and purified human islets were obtained from 5 non-diabetic donors from NIDDK-funded Integrated Islet Distribution Program (IIDP) at City of Hope (3 donors) and Prodo Laboratories, Inc. (2 donors) for intracellular GABA content experiments. Human islets were removed from the shipping container, centrifuged at 180 x g for 2 minutes, and placed into Prodo Islet Media, PIM (S), (PRODO Labs PIM-S001GMP) supplemented with PIM(ABS) (Prodo Labs PIM-ABS001GMP), PIM(G) (Prodo Labs PIM-G001GMP), and PIM(3x) (Prodo Labs PIM-3X001GMP). Islets were separated into non-adherent cell culture dishes at a density of 1000 islet equivalents per dish. Dishes were cultured at 24 °C upon initial receipt and up to one-week post receipt.

### Rat Pancreatic Islet Isolation and Perifusion

All experimental protocols were approved by the University of Florida Animal Care and Use Committees. Rat islets were isolated from pancreases of male and female postnatal day 5 (P5) Sprague Dawley rats (Charles River). Rat pups were killed by decapitation and the whole pancreas was removed and digested in 0.15 mg ml^−1^ Liberase TL (Roche 05401020001) in Hank’s buffered salt solution (HBSS) (Gibco 24020091) with 20 mM HEPES for 7 min with strong manual agitation. The Liberase enzyme was stopped with addition of HBSS, 20 mM HEPES and 0.5% FBS. Digested pancreas tissue was washed four times with HBSS, 20 mM HEPES and 0.5% FBS at 4 °C, resuspended in Histopaque-1119 (Sigma no. 11191), overlain with HBSS, 20 mM HEPES and 0.5% FBS (room temperature), and centrifuged for 20 min at 300*g* at 20 °C. Islets were collected from the interface between Histopaque and HBSS phases and washed three times with HBSS, 20 mM HEPES and 0.5% Newborn calf serum (NBCS). Islets were then hand-cleaned with a 200-µl pipette and cultured in 10-cm nonadherence petri dishes at 37 °C and 5% CO_2_, with 500 islets per dish, and 9 ml RPMI 1640 medium with GlutaMAX (Gibco no. 61870010), 10% FBS and 1% penicillin/streptomycin.

Isolated islets were used in conjunction with the Biorep Perifusion v2.0 system. Chambers loaded with a combination of 25 islet equivalents and Bio-Gel P-4 gel (Bio-Rad 150124) were loaded into the machine and perfused with KRBH buffer at a rate of 100 µL per minute. The protocol started with a 60-min equilibration period using heated 3 mM glucose KRBH, followed by a collection period of 10 min of 3 mM glucose, 20 min of 16.7 mM glucose, 15 min of 3 mM glucose, 5 min of 30 mM KCl, and 20 min 3 mM glucose. Perfusate was collected using a fraction collector every 2 minutes.

### KRBH Buffer Preparation

Krebs-Ringer Bicarbonate-HEPES Buffer (KRBH) was prepared with 115 mM NaCl, 4.7 mM KCl, 2.5 mM CaCl_2_, 1.2 mM KH_2_PO_4_, 1.2 mM MgSO_4_, 25 mM NaHCO_3_, 25 mM HEPES, and 0.2% BSA. Varying mM concentrations of glucose were added (1, 3, 5, 11, 16.7 mM) and indicated by the number and G before the buffer. 30 mM KCl was also added for select solutions and indicated as 30 mM KCl KRBH. Hypotonic KRBH was made with a lower NaCl concentration and included 90 mM NaCl, 4.7 mM KCl, 2.5 mM CaCl_2_, 1.2 mM KH_2_PO_4_, 1.2 mM MgSO_4_, 25 mM NaHCO_3_, 25 mM HEPES, and 0.2% BSA.

### Static Incubation of Islets

Groups of 100 human islets were handpicked on the Leica M80 stereomicroscope into 1 mL of 1G KRBH. Islets were pre-incubated for 1 hour at 37°C. Islets were moved to 250 µL 3G KRBH for 1 hour at 37°C. Islets were moved to 250 µL 16.7G KRBH for 1 hour at 37°C. Remaining solution from 3G and 16.7G incubation was collected, spun down at 5000 x g for 2 minutes, and collected into a microcentrifuge tube. Supernatant was frozen at –80 °C until analysis. Islets were moved to 1mL PBS. Islets were spun down at 800 x g for 3 minutes. The supernatant was removed. 200 µL RIPA buffer was added, and islets were sonicated for 10 seconds at 20% power (QSonica CL-334). Vehicle control (DMSO for diazoxide, ethanol for forskolin), 125 µM diazoxide (Sigma D9035), or 10 µM forskolin (Tocris 1099) were added into the 16.7G KRBH incubation.

Mouse islets were also collected in groups of 100 after isolation. Pre-incubation was done in 0G KRBH for 30 minutes. Incubations went from 5G KRBH to 11G KRBH. No other changes from the human islet protocol were observed. Short static incubation of mouse islets involved 50 islets in 100 µL of media. Islets were pre-equilibrated in 5G and then statically incubated for only 10 minutes in 5G and 11G.

Intracellular GABA was taken from 10 islets incubated in either low or high glucose solutions for 30 minutes. Islets were lysed and analyzed as described for HPLC.

Insulin secretion was analyzed using human insulin ELISA from ALPCO (80-INSHU-CH10) and rodent insulin ELISA from ALPCO (80-INSMR-CH10).

### HPLC for Whole Islets and Secreted GABA

GABA, taurine, and glutamate standards were made of 10 µM, 1 µM, 100 nM, and 10 nM in 1/3 1G KRBH + 0.05% Glutamax and 2/3 methanol. Samples from static incubations were diluted 1:2 in methanol. Whole islets were lysed in mobile phase containing 7% acetonitrile and 13% methanol in a phosphate buffer, using vigorous agitation. Lysate was then diluted 1:2 in 100% methanol. Standards, diluted samples, and 4 mM 2-mercaptoethanol/o-pthalaldehyde (OPA) were filtered through a 0.22 µm cellulose acetate membrane (Corning 8160) in a centrifuge at 5000g for 2 minutes. 20 µL of the standards (3 replicates) and samples (2 replicates) were loaded into a 96-well detection plate (Thermo Fisher AB-1400). 20 µL of OPA were loaded into a corresponding 96-well detection plate with the same number of wells filled as the sample/standards plate. Plates were sealed (Nacalai R37.001.00). The Eicom HTEC-510 HPLC with Insight AS-700 autosampler with FA-3ODS separation column was used to analyze the samples and standards (Sup. Fig 1A). A standard curve for GABA was created using a quadratic fit curve (Sup. Fig 1B). The standard curve was used to interpolate values of each sample’s GABA content from the sample chromatograms (Sup. Fig 1C).

**Figure 1:**
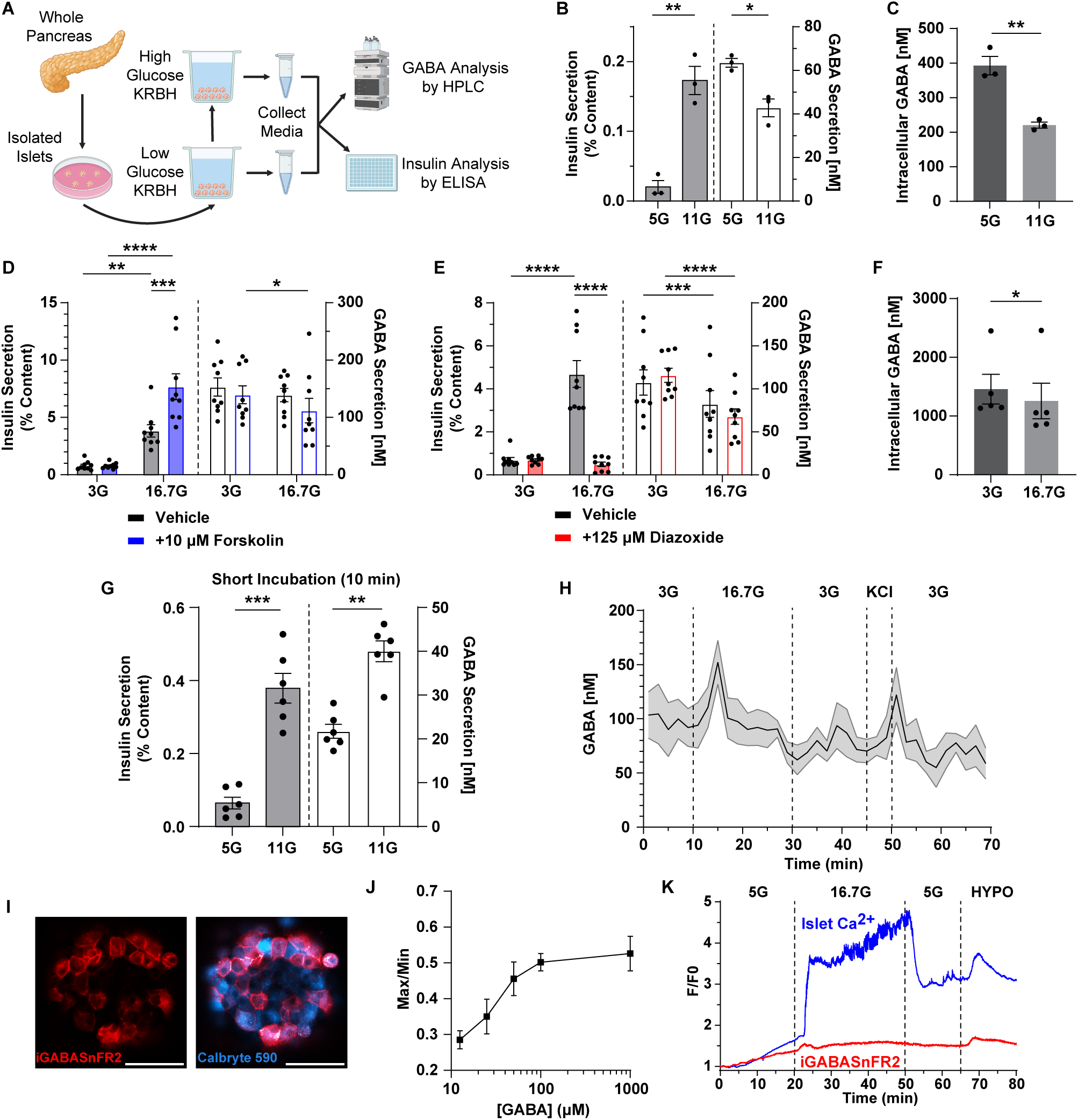
GABA Is Not Co-Regulated with Insulin. (**A**) Schematic of static incubations. Pancreatic islets are isolated from whole pancreas. Islets are stimulated in low glucose, then high glucose, and supernatants are collected and analyzed by HPLC and ELISA for GABA and insulin respectively. Created in https://BioRender.com. **(B-C)** C57BL/6J mouse islets incubated for 1 hour in 5 mM glucose and 11 mM glucose. **(B)** Insulin secretion and GABA secretion (n = 3, paired Student’s t-test, * p < 0.05, ** p < 0.01). **(C)** Intracellular GABA content from 10 C57BL6/J islets incubated in 5 mM and 11 mM glucose for 30 minutes (n = 3, unpaired Student’s t-test, ** p < 0.01). **(D)** Insulin and GABA secretion from static incubation of human donor islets incubated in 3 mM glucose and 16.7 mM glucose + vehicle or 10 μM forskolin for one hour (n = 3 donors, 3 replicates per donor, two-way matched ANOVA with Tukey’s post-hoc test, * p < 0.05, ** p < 0.01, *** p < 0.001, **** p < 0.0001). **(E)** Insulin and GABA secretion from static incubation of human donor islets incubated in 3 mM glucose and 16.7 mM glucose + vehicle or 125 μM diazoxide (n = 3 donors, 3 replicates per donor, two-way matched ANOVA with Tukey’s post-hoc test, *** p < 0.001, **** p < 0.0001). **(F)** Intracellular GABA content from 10 human islets incubated in 3 mM and 16.7 mM glucose for 30 minutes (n = 5 donors, unpaired Student’s t-test, * p < 0.05). **(G)** Insulin and GABA Secretion of C57BL/6J mouse islets incubated for 10 minutes in 5 mM glucose and 11 mM glucose (n = 6, paired Student’s t-test, ** p < 0.01, *** p < 0.001). **(H)** Neonatal Sprague Dawley rat islet perifusion measured by HPLC (± SEM, n = 3). **(I)** iGABASnFR2 expressed in C57BL/6J islets loaded with Calbryte 590 (Scale bar = 50 μm). **(J)** Titration curve of iGABASnFR2 plotted as maximum peak height over baseline divided by baseline (± SEM, n = 3). **(K)** Represenatative trace of iGABASnFR2 response to 16.7 mM glucose and hypotonic stimulation.

### GABA Biosensor Cells

GABA biosensor cells were obtained from Klemens Kaupmann at the Novartis Institutes for Biomedical Research in Basel, Switzerland. GABA biosensor cells consisted of CHO cells stably expressing heteromeric GABA_B_ receptors and the G-protein α subunit, Gαqo5, modified to couple to increases in [Ca^2+^]_i_ via InsP_3_^47, 48^. Biosensor cells were cultured in DMEM-F12 with 10% FBS, 2% geneticin, 1% glutamax, 0.4% hygromycin B, and 0.25% zeocin. Cells were removed from the culture plate with 0.25% trypsin-EDTA solution, enzymatic degradation was stopped with culture media, and spun down for 5 minutes at 180 x g. Cells were resuspended in KRBH with 5 mM glucose (5G). Biosensor cells were loaded with Calbryte 520 AM (AAT Bioquest 20651) for 30 minutes at 37°C. Islets were loaded with Calbryte 590 AM (AAT Bioquest 20701), or Calbryte 520 AM for tdTomato-expressing *Gad* ^βKO^ islets, in 5G KRBH for 30 minutes at 37°C.

A perfusion chamber (Warner RC-22) was adhered to a poly-L-lysine coated coverslip using vacuum grease. The chamber was filled with 200 µL of 5G KRBH, and the islets were placed into the chamber after rinsing the islets in 5G KRBH. The islets adhered to the coverslip in the chamber for 10 minutes before biosensor cells were added and left to rest for another 5 minutes. The perfusion chamber was imaged using confocal microscopy (Leica SP8) using an array of syringe pumps (New Era NE-1000) set to a 200 µL/min flow rate for each solution to be used, with multiple inlet solution lines converging into a single inlet via a manifold feeding into the perfusion chamber. A peristaltic pump (Ismatec Reglo ICC) was used to remove the waste solution from the perfusion chamber outlet.

Ca^2+^ traces for both islets and biosensor cells were normalized to F_0_, the average [Ca^2+^]_i_ intensity over the first 2 minutes of the recording. Biosensor cells were selected manually in ImageJ for cells that responded appropriately to the hypotonic and 10 µM GABA control stimulations and did not excessively flash regardless of solution (Sup. Fig 2). Cells far from the islets only responded to the 10 µM GABA control stimulation.

**Figure 2:**
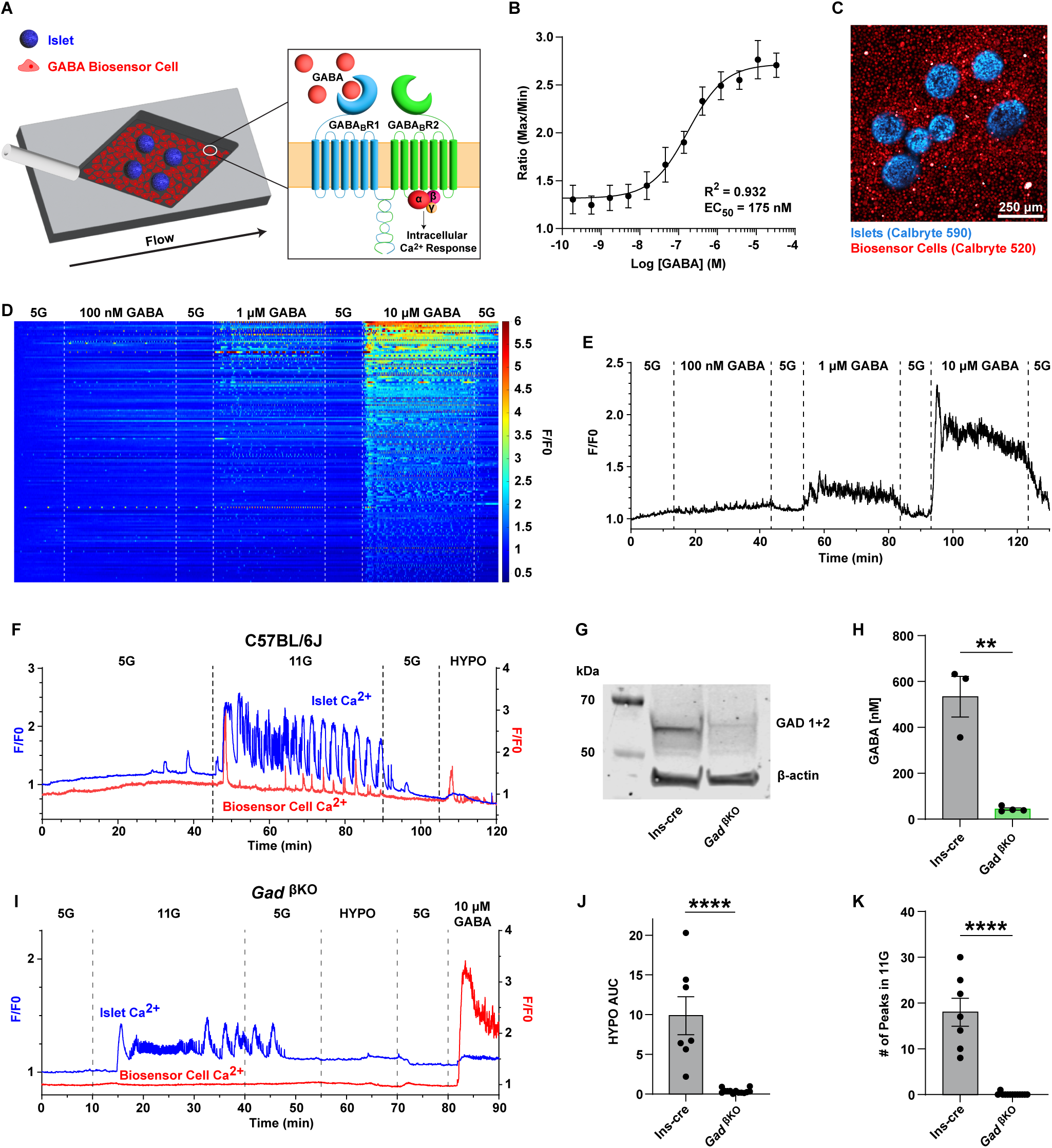
GABA Biosensor Cells Dynamically Detect GABA Release from Islets. (**A**) Schematic of GABA biosensor cell setup. In a microfluidic chip, islets are surrounded by GABA biosensor cells which will detect GABA release as solution flow over islets and cells. GABA biosensing is achieved by GABA_B_ receptor mediated calcium fluxes. **(B)** Log(agonist) vs. response curve for FLIPR assay performed with Calbryte 520 AM calcium dye for GABA biosensor cells (± SD, n = 16). **(C)** Representative image of islets with surrounding GABA biosensor cells. **(D)** Heatmap of individual GABA biosensor cell responses to increasing concentrations of GABA in 5 mM glucose KRBH. **(E)** Average trace of all individual biosensor cells in (D). **(F)** Representative calcium traces for single C57BL/6J islet and average surrounding biosensor cell calcium responses. **(G)** Western blot of Ins-cre and *Gad* ^βKO^ islets for GAD 1 and 2. **(H)** HPLC analysis of intracellular GABA content from Ins-cre (n = 3) and *Gad* ^βKO^(n = 4) islets (± SEM, student’s unpaired t-test, ** p < 0.01). **(I)** Representative calcium traces for single *Gad* ^βKO^ islet and average surrounding biosensor cell calcium responses. **(J)** Area under curve calculation of biosensor cell response for hypotonic stimulation (55 – 74 min) for Ins-cre (n = 7) and *Gad* ^βKO^(n = 13) islets (± SEM, student’s unpaired t-test, **** p < 0.0001). **(K)** Count of number of peaks during 11 mM glucose stimulation (10 – 40 min) for biosensor cells near Ins-cre (n = 7) and *Gad* ^βKO^(n = 13) islets (± SEM, student’s unpaired t-test, **** p < 0.0001).

### Matlab Analysis for HYPO AUC, Number of Peaks, Peak Alignment

Custom Matlab scripts were written to detect area under curve (AUC), number of peaks in islet and biosensor traces, and the alignment of the peaks between GABA biosensor cells and islet [Ca^2+^]_i_ oscillations.

### iGABASnFR2 Experiments

iGABASnFR2 constructs^49^ were put into adenoviral vectors with the rat insulin promoter and made into adenovirus by VectorBuilder (VB220519-1496ysk). Mouse C57BL6/J islets were isolated and transduced with virus for 24 hours. Islets were allowed to rest for another 24 hours in fresh media (RPMI 1650, 10% FBS, 1% Pen/Strep, 1% Glutamax) before imaging to ensure maximal sensor expression.

A perfusion chamber (Warner RC-22) was adhered to a poly-L-lysine coated coverslip using vacuum grease. The chamber was filled with 200 µL of 5G KRBH, and the islets were placed into the chamber after rinsing the islets in 5G KRBH. The islets adhered to the coverslip in the chamber for 10 minutes. The perfusion chamber was imaged using confocal microscopy (Leica SP8) using an array of syringe pumps (New Era NE-1000) set to a 200 µL/min flow rate for each solution to be used, with multiple inlet solution lines converging into a single inlet via a manifold feeding into the perfusion chamber. A peristaltic pump (Ismatec Reglo ICC) was used to remove the waste solution from the perfusion chamber outlet. Ca^2+^ traces for both islets and iGABASnFR were normalized to F_0_, the average [Ca^2+^]_i_ intensity over the first 2 minutes of the recording.

### Western Blot

Islet lysates were prepared by extraction of whole islet proteins in RIPA buffer (Thermo Fisher 89900) and a protease inhibitor (Roche 04693159001). Electrophoresis of proteins from islet lysates was performed using the Bolt western blot system. 40 µg of protein from each lysate were loaded into Bolt 4-12% Bis-Tris gels (Life Technolgies NW04120BOX) and run at a constant 200V for 25 minutes. Transfer was performed using standard transfer settings on the iBlot 2.0 device (Life Technologies). Membranes were blocked with 5% nonfat milk in tris buffered saline and incubated with primary antibodies overnight at 4 °C including custom rabbit anti-GAD65/GAD67 primary antibody^2, 50^ (1:1000) and mouse anti-β actin primary antibody (1:10000) (Sigma A1978). Membranes were then rinsed 3x with 1x PBS for 5 minutes each, then incubated in IRDye 680RD goat anti-rabbit secondary antibody (1:10000) (LICOR 922-68071) and IRDye 800CW goat anti-mouse secondary antibody (1:10000) (LICOR 922-68071) for one hour at room temperature. Membranes were then rinsed 3x with 1x PBS for 5 minutes each. Samples were rinsed again 3x in 1x PBS, followed by imaging on the LI-COR Odyssey CLx scanner.

### Adenovirus Transduction of Mouse Islets

Human Ad5 adenovirus was produced from plasmids for scramble shRNA (VectorBuilder VB010000-0021kwc) and LRRC8D shRNA (VectorBuilder VB240930-1432nab), both with mCherry reporters. C57BL/6J islets were transduced with virus overnight in RPMI 1640 with 10% FBS, and 1% pen/strep at 37 °C. The next day, islets were removed from viral media, rinsed in PBS, and replated in RPMI 1640 with 10% FBS, and 1% pen/strep at 37 °C.

### RT-qPCR

Human Ad5 adenovirus was produced from plasmids for scramble shRNA (VectorBuilder VB010000-0021kwc) [CCTAAGGTTAAGTCGCCCTCG] and LRRC8D shRNA (VectorBuilder VB900143-9202rsm) [GATGCTCACTTGCAAGTTATT] both with mCherry reporters. Mouse insulinoma 6 (MIN6) cells were plated at 100K/cm^2^ in a 12-well culture dish in high glucose DMEM with 15% FBS, 1% pen/strep, and 1% glutamax. The following day, viruses were added to the cells and left to rest overnight before removing viral media, rinsing cells with PBS, and replacing the media with fresh culture media. Cells were allowed to rest for another 24 hours before RT-qPCR was performed.

RNA was isolated using the Monarch Total Miniprep Kit (New England Biolabs T2010). RNA concentrations were measured using the NanoDrop 2000 (Thermo Fisher). Reverse transcriptase was performed with 1 µg RNA per sample with the LunaScript RT Supermix Kit (New England Biolabs E3010) on a Bio-Rad DNA Engine thermal cycler. qPCR was performed with Luna Universal qPCR Master Mix (New England Biolabs M3003) on a QuantStudio 6 Flex (Applied Biosystems). Primers for mouse LRRC8D (Qiagen QT02255085) and a control primer for mouse GAPDH (Qiagen QT01658692) were used.

### MIN6 Hypotonic Stimulation

Human Ad5 adenovirus was produced from plasmids for scramble shRNA (VectorBuilder VB010000-0021kwc) and LRRC8D shRNA (VectorBuilder VB240930-1432nab) both with mCherry reporters. Mouse insulinoma 6 (MIN6) cells were plated at 100K/cm^2^ in a 6-well culture dish in high glucose DMEM with 15% FBS, 1% pen/strep, and 1% glutamax. The following day, viruses were added to the cells and let rest for 6 hours before removing viral media, rinsing cells with PBS, and replacing with fresh culture media. Cells were allowed to sit overnight. Cells were then transfected with the human GAD65-GFP plasmid and then left to rest for an additional 48 hours before stimulation was performed.

MIN6 cells were then treated with 0.25% trypsin-EDTA to remove them from the well. Cells were spun down at 180 x g for 5 minutes and resuspended in 3G isotonic KRBH. Cells were then split in half into an isotonic group and a hypotonic group. Cells were again spun down and then resuspended in either 3G KRBH isotonic solution or 3G KRBH hypotonic solution. Cells were incubated at 37°C for 30 minutes. Cells were spun down again, and supernatant was collected and analyzed for GABA release via HPLC as described.

### Ca^2+^ FLIPR Assay

The FLIPR assay was completed as had been previously published^51^. CHO biosensor cells were cultured in DMEM-F12 media supplemented with 10% heat-inactivated FBS, 2% Geneticin, 1% Glutamax, 0.4% Hygromycin B, and 0.25% Zeocin. On the day of the assay, cells were dissociated with TrypLE, counted and seeded at 1,500 cells/well in 1536 well Aurora clear bottom low-based TC treated black plates in 3 µl KRBH 3G buffer with 1% DMSO. The plates were then treated with Calbryte 520 AM in Pluronic F-127 in 3G KRBH buffer in the 3 µl. The plates were incubated for 1 hour at 37°C in the dark. Measurements of fluorescence intensity (λ_ex_=470 nm, λ_em_=535nm) were made using a FLIPR Tetra. A read of basal fluorescence values was made (5 s) prior to addition of the GABA diluted in 75% DMSO by pintool (30 nl). 3 minutes of reads were collected. The raw data collected was the maximum for the 3-minute read divided by the average of the basal read. This ratio was used for the dose response curves.

### Vgat-ires-cre x Ai14 tdTomato Mouse

Vgat-ires-cre x Ai14 tdTomato mice were obtained from Dr. Sunil Gandhi’s lab at the University of California, Irvine^52, 53^. Fixed brain and pancreas were obtained from 4% paraformaldehyde (PFA) in PBS perfusion fixed mice.

### Immunostaining Mouse and Brain Sections

Mouse tissues were cryoprotected in 15% sucrose in PBS overnight at 4°C and then in 30% sucrose in PBS overnight at 4°C on consecutive days. Tissues were placed in OCT (Fisher 23-730-571) in cryomolds (Electron Microscopy Sciences 62642-01) and refrigerated before freezing tissues over isopentane (Sigma M32631) cooled by liquid nitrogen. Sections were made of the tissues using the Leica CryoStat CM1950.

For staining, slides were rinsed 2x with 1x PBS containing 0.3% TritonX (Sigma T9284) (TxPBS) for 5 minutes each. Tissues were blocked with 10% donkey serum in TxPBS for 1 hour. Primary antibodies were added in 1% donkey serum in TxPBS overnight at 4 °C. Tissues were rinsed 3x with TxPBS for 5 minutes each. Secondaries were added at room temperature in TxPBS for 1 hour. Tissues were rinsed 3x with TxPBS for 5 minutes each. Tissues were mounted to coverslips with Prolong Gold containing DAPI (Thermo Fisher P36935) and left to rest overnight at room temperature before imaging.

Antibodies Used:

Primaries:

VGAT: polyclonal rabbit, oyster 650 labeled, 1:1000, (Synaptic Systems 131103C5)

Insulin: polyclonal guinea pig, 1:1000 (DAKO A0564)

GABA: polyclonal rabbit, 1:1000 (Sigma A2052)

Secondaries:

Donkey anti-guinea pig CF488A: 1:200 (Sigma SAB4600033)

Donkey anti-rabbit Alex Fluor 647: 1:200 (Invitrogen A31573)

### Gene expression analysis from publicly available islet scRNA-seq datasets

Human and mouse islet scRNA-seq datasets were used to investigate the single-cell gene expression of genes of interest. Mouse islet scRNA-seq data from three C57BLKS wild type mice (wt/wt) were downloaded as Cell Ranger output files from the Gene Expression Omnibus (GEO) database, accession number: GSE200531^54^. Human islet scRNA-seq data from 65 Human Pancreas Analysis Program (HPAP) donors were downloaded as a Seurat object from http://www.isletgenomics.org/^55^. Both mouse and human datasets were loaded and analyzed in R Studio v2024.04.0 with the R package Seurat v5.1.0^56^.

Mouse islet scRNA-seq datasets were processed as follows: Seurat objects were created, and quality control was performed with visualization tools of Seurat. Seurat objects were filtered for < 7.5% mitochondrial reads and a minimum of 1000 to 7500 genes per cell. Doublets were removed with DoubletFinder. The Seurat object was then normalized and scaled. The 2000 most variable features were used for PCA dimensionality reduction with 20 principal components. Harmony v1.2.3^57^ was used to perform batch correction and for uniform manifold approximation and projection (UMAP). Clustering of cells was performed with the Leiden algorithm at a resolution of 0.5. Islet cell type genes were used to identify clusters and/or perform additional subclusters with 0.25 resolution. Clusters with residual doublets were manually removed if they expressed multiple cell type markers. Harmony algorithm was rerun, and final clusters were annotated based on their cell type marker expression.

Final Seurat objects from human and mouse scRNAseq datasets were filtered for endocrine cells. In addition, only HPAP donors with no diabetes were used for the gene expression analysis described here. Single-cell gene expression for each gene of interest was visualized as bar plots of log-scaled normalized expression data. Bar plots were created with ggbarplot function of ggpubr v0.6.0 (https://rpkgs.datanovia.com/ggpubr/)^58^ and patchwork v1.3.0 (https://patchwork.data-imaginist.com/)^59^. Heatmaps of beta cell gene expression across disease state were generated with the pseudo-bulk matrices for the beta cell cluster and visualized with ComplexHeatmap v2.22.0^60, 61^.

### Gene and protein expression analysis from tissues in the Human Protein Atlas database

The protein expression profiles of 15302 proteins based on immunohistochemistry (IHC) in 45 human tissues and the tissue annotation table were downloaded as tsv files from (https://www.proteinatlas.org/humanproteome/tissue/data#normal_tissues_ihc)^62^. Both files were loaded in R Studio v2024.04.0 for data visualization. The protein expression level for each protein was obtained for the following levels: “Not detected” (ND), “Low” (L), “Medium” (M), and “High” (H). In addition, the protein expression levels labeled as “Uncertain” reliability were excluded from further analysis. The tissue annotation table included the organ from which the tissue originated and was used to filter for organs of interest. The tissue: retina was manually annotated as organ: brain. The protein expression table was then filtered for organs: brain and pancreas, and cell types: neuronal cells (brain tissues), pancreatic endocrine cells and exocrine glandular cells (pancreas tissue). Bar plots of protein expression level (ND, L, M, H) were created with the graphical tools of ggpubr and patchwork and included tissue and cell type corresponding to each organ, color-coded by protein.

The consensus RNA expression data from 50 tissues based on transcriptomics data from both the Human Protein Atlas (HPA) and GTEx were downloaded as tsv files from https://www.proteinatlas.org/humanproteome/tissue/data#consensus_tissues_rna, including the tissue annotation table. The consensus transcriptomics data from HPA shows the gene expression level as normalized transcripts per million (nTPM), which was calculated as the maximum nTPM value for each gene in the two data sets. Similarly, if multiple sub-tissues are present, the maximum value was considered. The tissue: retina was manually annotated as organ: brain. Similarly to protein expression, the tissue RNA expression was obtained for the organs: brain and pancreas. Gene expression for each gene of interest is presented as bar plots per tissue, color-coded by gene.

### Statistical Analysis

Statistical analysis was performed in GraphPad Prism 10. Analysis of variance (ANOVA) was used when comparing between multiple groups while student’s unpaired t-test was used when only comparing two groups. Tukey’s post-hoc test was used to make individual comparisons in ANOVA tests. Specific test, significance values, and error bars are detailed in each figure caption for each graph.

## Results

### GABA is not co-regulated with insulin

Previous studies conflict as to whether GABA is co-released with insulin during high glucose stimulation^6, 24, 25^. To investigate GABA co-secretion with insulin, C57BL/6J mouse islets were statically incubated for one hour in low glucose followed by one hour in high glucose. GABA release to the supernatant was measured by HPLC with electrochemical detection (Sup. Figure 1), and insulin release was measured by ELISA from the same samples (Figure 1A). Insulin release increased from low glucose to high glucose while GABA release slightly decreased from low to high glucose (Figure 1B). Intracellular GABA content also decreased from low to high glucose, following the GABA secretion (Figure 1C). These results indicate that cumulative GABA release from mouse islets does not change in the same manner as insulin secretion does and is even reduced during long incubations in high glucose.

To further test GABA’s co-release with insulin, human islets from three different human donors were obtained. We used forskolin and diazoxide to stimulate or inhibit insulin secretion, respectively, during one-hour static glucose stimulation. Forskolin doubled insulin secretion in high glucose, but the GABA secretion was unchanged compared to the vehicle control (Figure 1D). Diazoxide ablated insulin secretion in high glucose, but again, the GABA secretion was unchanged compared to the vehicle control (Figure 1E). Intracellular GABA content followed the GABA secretion trend, showing a significant decrease from low to high glucose in human islets (Figure 1F). C57BL/6J mouse islets showed a similar decrease in both secreted and intracellular GABA when stimulated with high glucose. Cumulative GABA secretion from mouse and human islets over a one hour static incubation is not sensitive to factors that modulate insulin secretion, in agreement with our previous results that GABA secretion not regulated by glucose^9^.

We next sought to distinguish between different time scales that may affect the dynamics GABA release. C57BL/6J islets were statically incubated in low and high glucose for ten minutes to capture just the initial glucose response period. In short static incubations, GABA and insulin release was significantly higher in high glucose (Figure 1G). Because glucose stimulated GABA release during the acute post-stimulate phase but not over longer time periods, we also analyzed GABA secretion dynamically during islet perifusion. To detect GABA by HPLC in perifusate, we used rat islets as greater numbers of islets were needed due to the higher dilution factor of the collected fractions. When examined dynamically, there was an acute pulse of GABA release immediately upon high glucose or KCl stimulation (Figure 1H) that did not persist during continued glucose stimulation. Perifusion with HPLC has relatively poor temporal resolution (1-2 min per time fraction) and requires large numbers of islets to reach the threshold of detection. We next explored optical biosensor methods to measure GABA release in real time from single islets with better temporal sensitivity.

We measured dynamic GABA secretion from single islets using a fluorescent GABA-sensing genetically encoded reporter, iGABASnFR2. iGABASnFR2 detects GABA binding directly via increases in cpGFP fluorescence upon GABA binding^49^. We transduced C57BL6/J mouse islets with an adenovirus encoding iGABASnFR2 (Figure 1I) and imaged these islets under varying GABA concentrations to create a titration curve for activity (Figure 1J). We observed iGABASnFR2 sensitivity to applied GABA in the 10-100 µM range. This is relatively consistent with the published EC_50_ of iGABASnFR2 of 6.4 µM, determined in neurons^49^. Using this genetically encoded biosensor combined with simultaneous measurement of [Ca^2+^]_i_ with Calbryte 590 AM, GABA release was observed during the acute [Ca^2+^]_i_ increase in response to high glucose but not during continued high glucose stimulation. Interestingly, iGABASnFR2 did not detect phasic secretion events akin to synaptic GABA release from neurons^49^ or insulin granule exocytosis events^63^ (Figure 1K). Instead, the GABA secretion event was uniform throughout the islet, indicative of tonic secretion. To further test this concept, we triggered opening of VRAC with hypotonic stimulation which results in tonic GABA secretion from the cytosol. Hypotonic stimulation generated a GABA release event that was similar to acute high glucose (Figure 1K). This result confirms the presence of an initial burst of GABA release during the first phase [Ca^2+^]_i_ response of beta cells, but that GABA is not continuously released at a high level during continued glucose stimulation. While iGABASnFR produced excellent temporal resolution, it also showed limited sensitivity to detect changes in the extracellular GABA concentration in the islet. To obtain better concentration sensitivity in dynamic GABA secretion experiments, we next implemented a biosensor cell system that incorporates signal amplification.

### GABA biosensor cells dynamically detect GABA release from islets

To provide highly sensitive temporal and concentration dynamics of GABA release, we implemented a real-time biosensor cell assay using CHO cells expressing GABA_B_R coupled to a modified G-protein α subunit, Gαqo5, which is coupled to [Ca^2+^]_i_ via InsP3 (Figure 2A). Using a fluorescence imaging plate reader (FLIPR) assay for [Ca^2+^]_i_, we measured the GABA sensitivity of the biosensor cells to have an EC_50_ = 175 nM (Figure 2B) and dynamic range of approximately 10 nM to 10 µM. To measure GABA secretion dynamics, we seeded mouse islets in an open bath flow chamber surrounded by a dense lawn of the GABA biosensor cells. We then flowed different solutions across the islets using a custom syringe pump array and monitored [Ca^2+^]_i_ in the biosensor cells and islets simultaneously by loading each cell type with a different Ca^2+^ indicator Calbryte 520 AM and Calbryte 590 AM, respectively, and imaging live by confocal (Figure 2C). The biosensor cells respond to increasing GABA concentrations with increasing [Ca^2+^]_i_ (Figure 2D). As individual biosensor cells varied somewhat in sensitivity to GABA, the average response of several responding cells in close proximity to an islet was used to represent the total response (Figure 2E, Sup. Fig 2).

Dynamic glucose stimulation was performed with C57BL/6J islets adjacent to GABA biosensor cells to investigate GABA secretion. The islet [Ca^2+^]_i_ trace was quiet during low glucose (5G) stimulation, but the islet responds and begins to oscillate following high glucose (11G) stimulation. The biosensor cells detected a large initial release of GABA immediately following high glucose. This large initial release was followed by periodic pulses of GABA secretion. As a positive control for islet GABA secretion, we used hypotonic stimulation, which swells the islet and activates VRAC, causing GABA release from the cytosol. Hypotonic stimulation resulted in another large biosensor cell detection of GABA release (Figure 2F). The strongest GABA release peaks were detected during the islet first-phase Ca^2+^ response and during the hypotonic stimulation.

To confirm the biosensor cells respond specifically to only GABA and not to other co-released molecules, we generated a knockout mouse that lacks GABA biosynthesis in beta cells (*Gad* ^βKO^) by crossing mice floxed for *Gad1* and *Gad2* with Ins-Cre. *Gad* ^βKO^ islets had no GAD expression by western blot (Figure 2G) and were devoid of GABA by HPLC (Figure 2H), indicating a successful knockout compared to the Ins-cre control. Biosensor cells were silent in proximity to *Gad* ^βKO^ islets including during high glucose and hypotonic stimulation (Figure 2I-K, Sup. Fig 3) compared to Ins-cre control (Sup. Fig 4). This result indicates the biosensor cells responded specifically to GABA and did not respond to other co-released molecules. The simultaneously measured islet Ca^2+^ response of *Gad* ^βKO^ islets differed from that of C57BL/6J or Ins-cre controls. We have separately published a detailed study characterizing the *Gad* ^βKO^ mice and the Ca^2+^ dynamics of their isolated islets^64^. To summarize, *Gad* ^βKO^ islets exhibit a long period of maximal activation following glucose stimulation, delayed initiation of Ca^2+^ oscillations, lower amplitude of Ca^2+^ oscillations, and insulin hyper-secretion. These *Gad* ^βKO^ Ca^2+^ signaling trends are apparent in the representative traces (Figure 2F, I). The difference in Ca^2+^ signaling likely results from the loss of GABA signaling, which we hypothesized to operate as a feedback mechanism that strengthens Ca^2+^ oscillations in islets.

**Figure 3:**
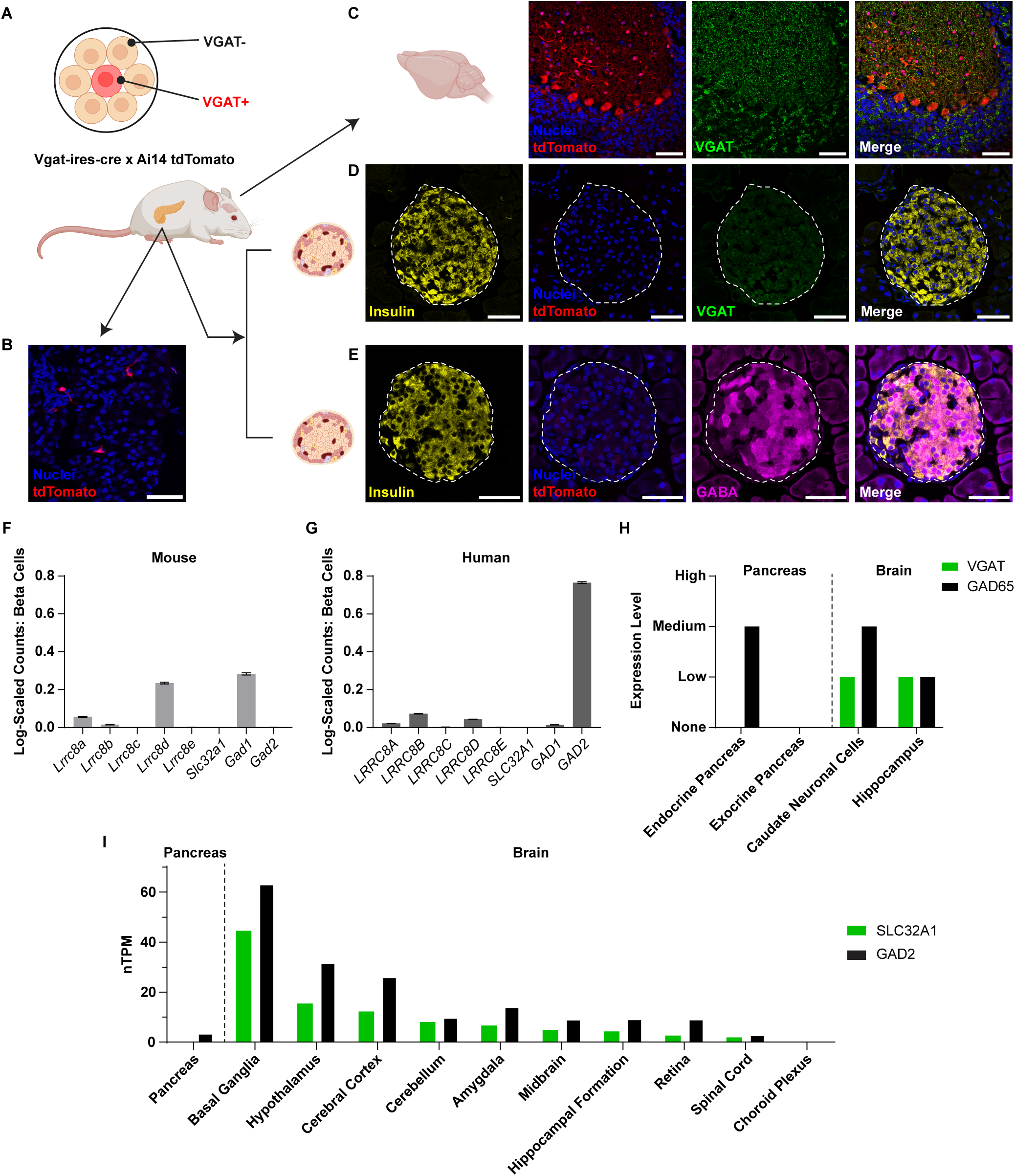
Vesicular GABA Transporter Is Not Present in Beta Cells. (**A**) Schematic of Vgat-ires-cre x Ai14 tdTomato reporter mouse. Created in https://BioRender.com. **(B)** Immunohistochemistry confocal images of reporter mouse pancreatic neuronal-like tdTomato+ structures (Scale bar = 50 μm). **(C)** Immunohistochemistry confocal images of reporter mouse hippocampal brain tissue stained for DAPI (nucleus) and vesicular GABA transporter (VGAT) with tdTomato labeled cells (Scale bar = 50 μm). **(D)** Immunohistochemistry confocal images of reporter mouse pancreatic islet lacking tdTomato-positive cells and VGAT (Scale bar = 50 μm). **(E)** Immunohistochemistry confocal images of reporter mouse pancreatic islet with GABA and without tdTomato (Scale bar = 50 μm). **(F)** Pancreatic endocrine single cell RNA-seq data of C57BLKS wild type mice for *Gad1* and *2*, *Slc32a1*, and *Lrrc8a-e*. **(G)** Pancreatic endocrine single cell RNA-seq data of donors without diabetes from HPAP for *GAD1* and *2*, *SLC32A1*, and *LRRC8A-E*. **(H-I)** Protein **(H)** and gene **(I)** expression of GAD65 (*GAD2*) and VGAT (*SLC32A1*) in humans by tissue type from the Human Protein Atlas.

**Figure 4:**
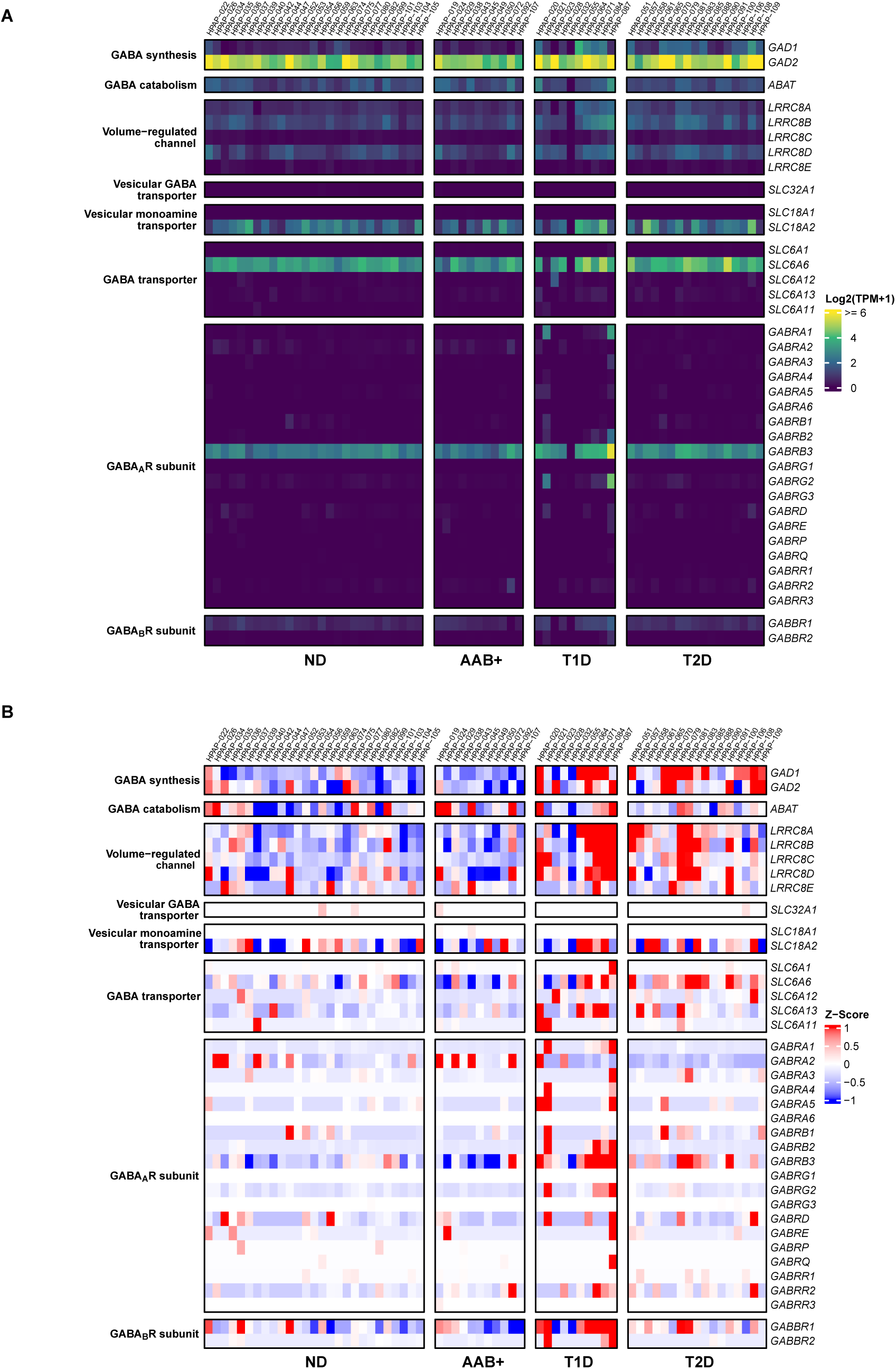
Pseudobulk Gene Expression from Single-cell RNA Sequencing of Human Beta Cells. Single-cell RNA sequencing of human beta cells from HPAP data set in various diabetes disease states shown as log-normalized expression **(A)** and differential expression as z-scores **(B)** (ND = no diabetes, AAB+ = autoantibody positive, T1D = type 1 diabetes, T2D = type 2 diabetes).

### Vesicular GABA transporter is not present in beta cells

We wanted to understand if the islet GABA secretion is from a vesicular or cytosolic source. We investigated whether beta cells contained the vesicular GABA transporter (VGAT) encoded by the *SLC32A1* gene, which transports GABA into vesicles in neurons and would be required for loading GABA into insulin granules or synaptic-like microvesicles^21^. Previous literature has demonstrated both evidence for and against VGAT expression in beta cells in rat, mouse, and human islets^9, 26, 27, 28, 29^. As VGAT antibody staining in pancreas has been inconsistent, we obtained pancreas and brain tissue from VGAT reporter mice, generated by crossing mice harboring a Cre-dependent fluorescent protein reporter (tdTomato) with mice expressing Cre-recombinase under the control of the VGAT promoter^52^. We investigated tdTomato expression alongside VGAT immunostaining in the brain and pancreas (Figure 3A). Sections of exocrine pancreas showed neuronal-like tdTomato+ structures (Figure 3B). Sections of hippocampus showed prominent tdTomato and VGAT expression (Figure 3C). The tdTomato and VGAT signals in neurons do not colocalize because tdTomato is a cytosolic reporter while VGAT is targeted to synaptic vesicles at axon terminals. In islets, tdTomato was entirely absent (Figure 3D-E). Consistent with some previous reports^26, 27, 28^, there was a faint immunoreactivity in islets for the VGAT antibody, which we interpret as non-specific given the lack of tdTomato expression and much brighter VGAT immunoreactivity in brain. We also immunostained pancreatic islets for GABA using an antibody we have previously validated. Interestingly, GABA content in mouse islets was heterogenous, showing that some beta cells are negative for GABA, reflecting our previous reports of heterogenous GAD/GABA expression in rodent islets^25^. Human islets have uniform GAD expression and GABA content^25^. It is currently unknown whether GAD/GABA positivity in mouse beta cells represents the different beta cell subtypes that help to polarize and coordinate whole islet Ca^2+^ dynamics.

We next investigated whether islet VGAT expression could be observed in a public scRNA-seq dataset of mouse islets^54^. Mouse beta cells were observed to have high expression of GAD67 (*Gad1*), but VGAT (*Slc32a1*) was below the detection threshold (Figure 3F). Similarly, in the human islet scRNA-seq data from donors with no diabetes in the HPAP dataset^55^, human beta cells showed high expression of GAD65 (*GAD2*) but undetectable VGAT (*SLC32A1*) (Figure 3G). The two GAD isoforms are known to be differentially expressed by species^2, 3^. In the Human Protein Atlas (HPA) dataset, VGAT protein and RNA were only detected in the brain and not in the pancreas tissue (Figure 3H-I)^62^. Barring the existence of an alternative vesicular GABA transporter, the absence of VGAT expression in the pancreas suggests that neuron-like vesicular release is unlikely to be the primary mechanism of GABA secretion from beta cells.

### Pseudobulk gene expression from single-cell RNA sequencing of human beta cells

While VGAT may not be present in beta cells, we also wanted to look at the expression of the full set of known components of the GABA system using the HPAP human islet single-cell dataset (Figure 4A)^55^. As expected, GAD65 (*GAD2*) is the dominant gene for human beta cell GABA synthesis. GABA transaminase (ABAT) is expressed for GABA catabolism. VRAC genes LRRC8A, B, and D are expressed, of which with LRRC8A/D isoform is permeable to GABA for efflux. Vesicular GABA transporter VGAT (SLC32A1) is not detected. A vesicular monoamine transporter, VMAT2, is detected, which may have low-affinity for GABA in vitro, but is shown to not transport GABA under physiological conditions^33^. For GABA re-uptake from the extracellular environment, we found that among the membrane GABA transporters only the taurine transporter (TauT, *SLC6A6*) is detected. TauT has the ability to transport GABA^65^, and has been shown to function for GABA uptake in islets, but not release^9^. The most highly expressed GABA_A_R gene encodes the GABA_A_R beta-3 subunit, followed by the alpha-2 subunit, consistent with previous reports^16^. Human islets also express GABA_B_R1 but GABA_B_R2 was not detected, consistent with previous reports^14, 16^. Because GABA_B_R is an obligate heterodimer, it has been suggested that human beta cells lack a functional GABA_B_R unless expression is induced^14^. We also looked for changes to the GABA system in different states of diabetes and observed few differences between donors with no diabetes and autoantibody positive donors (Figure 4B). However, GABA synthesis and VRAC gene expression was increased in both type 1 and 2 diabetes while GABA_A_R and GABA_B_R gene expression was increased in type 1 diabetes (Figure 4B). To summarize, single-cell RNA sequencing data from multiple donors gives an overview of the human islet GABA system that consists of GAD65 and GABA transaminase for synthesis and catabolism, LRRC8A/D for cytosolic efflux, functional signaling dominated by GABA_A_R beta-3, and TauT for re-uptake, but that lacks a mechanism for vesicular secretion.

### GABA is secreted via the LRRC8A/D channel

VRAC is a Cl^-^-permeable membrane channel that is activated upon cell swelling, causing a regulatory volume decrease and release of osmolytes from the cell, including GABA and taurine^36^ (Figure 5A). The VRAC channel is a hexamer made up of a combination of 5 leucine-rich repeat protein subunits, LRRC8A-E, with each channel requiring the LRRC8A subunit (also known as SWELL1) for its function^66^. The GABA-permeable VRAC isoform, LRRC8A/D, is expressed in mouse and human beta cells (Figures 3F-G, 4A). We previously confirmed a role for LRRC8A in beta cell GABA secretion^9^ but did not investigate LRRC8D.

**Figure 5:**
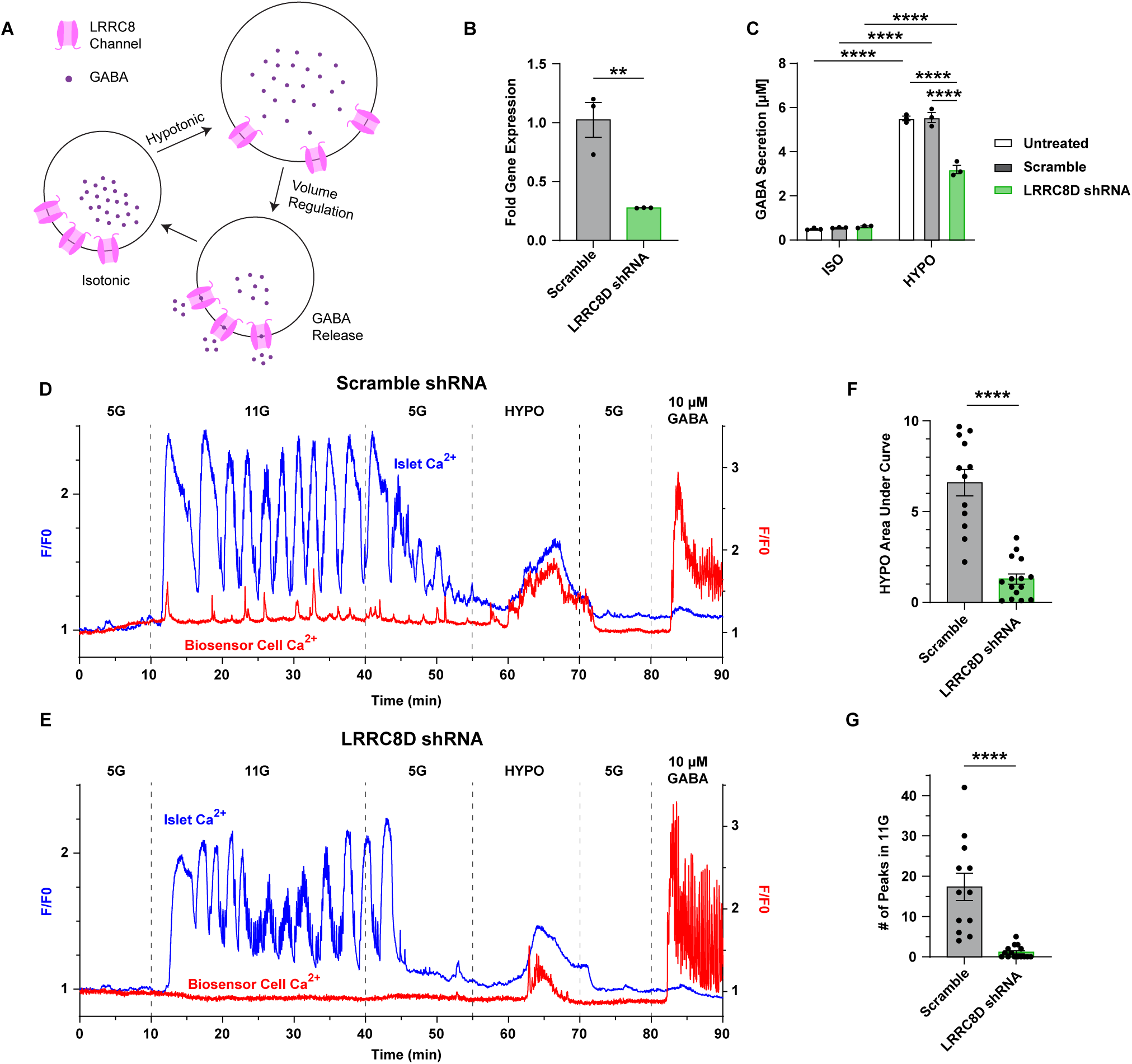
GABA Is Secreted via the LRRC8D Channel. Adenovirus containing scramble shRNA or LRRC8D shRNA was used to transduce mouse insulinoma 6 beta cells (MIN6) (B-C) and C57BL/6J islets (D-G). **(A)** Schematic of hypotonic stimulation on GABA release via LRRC8 channels in beta cells. **(B)** Fold gene expression of MIN6 cells transduced with scramble or LRRC8D shRNA (n = 3, ± SEM, student’s unpaired t-test, ** p < 0.01). **(C)** GABA secretion in isotonic or hypotonic solution of untreated, scramble shRNA transduced, and LRRC8D shRNA transduced MIN6 cells (n = 3, ± SEM, two-way ANOVA with Tukey’s post-hoc test, **** p < 0.0001). **(D)** Representative calcium trace of single C57BL/6J islet transduced with scramble shRNA with average surrounding biosensor cell calcium responses. **(E)** Representative calcium trace of single C57BL/6J islet transduced with LRRC8D shRNA with average surrounding biosensor cell calcium responses. **(F)** Area under curve calculation of biosensor cell response for hypotonic stimulation for scramble (n = 12) and LRRC8D shRNA (n = 15) transduced islets (± SEM, student’s unpaired t-test, **** p < 0.0001). **(G)** Count of number of peaks during 11 mM glucose stimulation for biosensor cells near scramble (n = 12) and LRRC8D shRNA (n = 15) transduced islets (± SEM, student’s unpaired t-test, **** p < 0.0001).

We generated an adenovirus encoding *Lrrc8d* shRNA to determine the role of LRRC8D in islet GABA secretion. The shRNA reduced *Lrrc8d* mRNA expression by 70% (Figure 5B) and decreased hypotonically-induced GABA secretion by 50% in MIN6 beta cells (Figure 5C). Next, we performed dynamic biosensor cell assays on C57BL/6J mouse islets with scramble and *Lrrc8d* shRNAs (Figure 5D-E, Sup. Figures 5-6) and observed that the *Lrrc8d* knockdown resulted in less GABA release in hypotonic stimulation and fewer GABA release events during high glucose (Figure 5F-G). This demonstrates a large role for LRRC8D mediating beta cell GABA release and specifically, GABA pulses.

**Figure 6:**
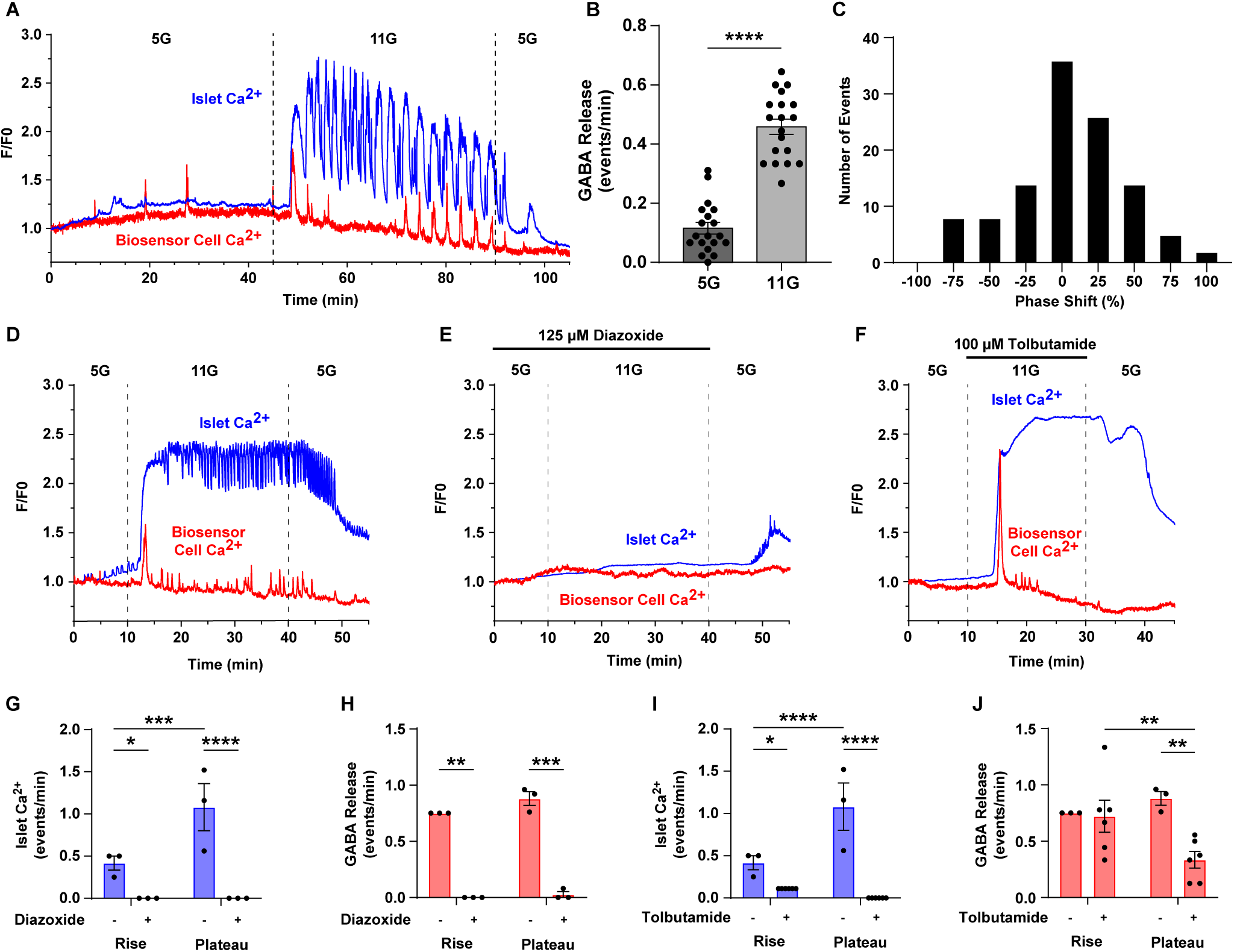
GABA Secretion Is Coordinated with [Ca^2+^]_i_ Increases in Beta Cells. (**A**) Representative trace of C57BL/6J islets on GABA biosensor cells with slow islet oscillations. **(B)** Quantification of GABA release events in low glucose (5G) and high glucose (11G) (± SEM, n = 19, paired student’s t-test, **** p < 0.0001). **(C)** Histogram of alignment between GABA biosensor cell peaks and islet Ca^2+^ peaks for islet oscillations > 1 minute in length. 0% shift indicates perfect alignment. **(D-F)** Representative calcium traces for single C57BL/6J islet and average surrounding biosensor cell calcium responses in **(D)** control conditions, **(E)** 125 μM diazoxide, and **(F)** 100 μM tolbutamide. **(G-H)** Calculation of islet calcium flux **(G)** and GABA release **(H)** for initial calcium rise (11 – 15min) and calcium plateau (15 – 40 min) for control and diazoxide-treated islets (n = 3, ± SEM, two-way matched ANOVA with Tukey’s post-hoc, * p < 0.05, ** p < 0.01, *** p < 0.001, **** p < 0.0001). **(I-J)** Calculation of islet calcium flux **(I)** and GABA release **(J)** for initial calcium rise (CTRL: 11 – 15min, Tol: 13 – 22 min) and calcium plateau (CTRL: 15 – 40 min, Tol: 22 – 30 min) for control (n = 3) and tolbutamide-treated (n = 6) islets (± SEM, two-way matched ANOVA with Tukey’s post-hoc, * p < 0.05, ** p < 0.01, **** p < 0.0001).

### GABA secretion is coordinated with [Ca^2+^]_i_ oscillations in beta cells

After observing the loss of high glucose GABA release events with *Lrrc8d* shRNA, we wanted to further investigate the timing and coordination of these release events. In addition, we were interested to understand if trends in glucose control over cumulative GABA release over long and short incubations observed by HPLC (Figure 1) would be similarly reflected in the biosensor cell assay. Thus, we performed long incubations of 45 minutes each in low and high glucose while monitoring GABA secretion by biosensor cells (Figure 6A-C, Sup. Figure 7). As before, we observed discrete GABA secretion pulses including a single large pulse immediately upon first-phase Ca^2+^ activation. Following, the initial activation period and start of deep periodic oscillations, GABA release events tended to be well-aligned with the peaks of individual Ca^2+^ waves. We also observed that the total number of GABA release events significantly increased during high glucose stimulation. Phase-shift analysis of GABA and Ca^2+^ peaks demonstrated the two signals and were predominantly in-phase (Figure 6A-C, Sup. Fig. 7). The GABA secretion events and islet [Ca^2+^]_i_ suggest that the mouse islet LRRC8A/D-dependent GABA release is coordinated with rhythmic activities that align with the timing of slow-wave [Ca^2+^]_i_ oscillations.

**Figure 7:**
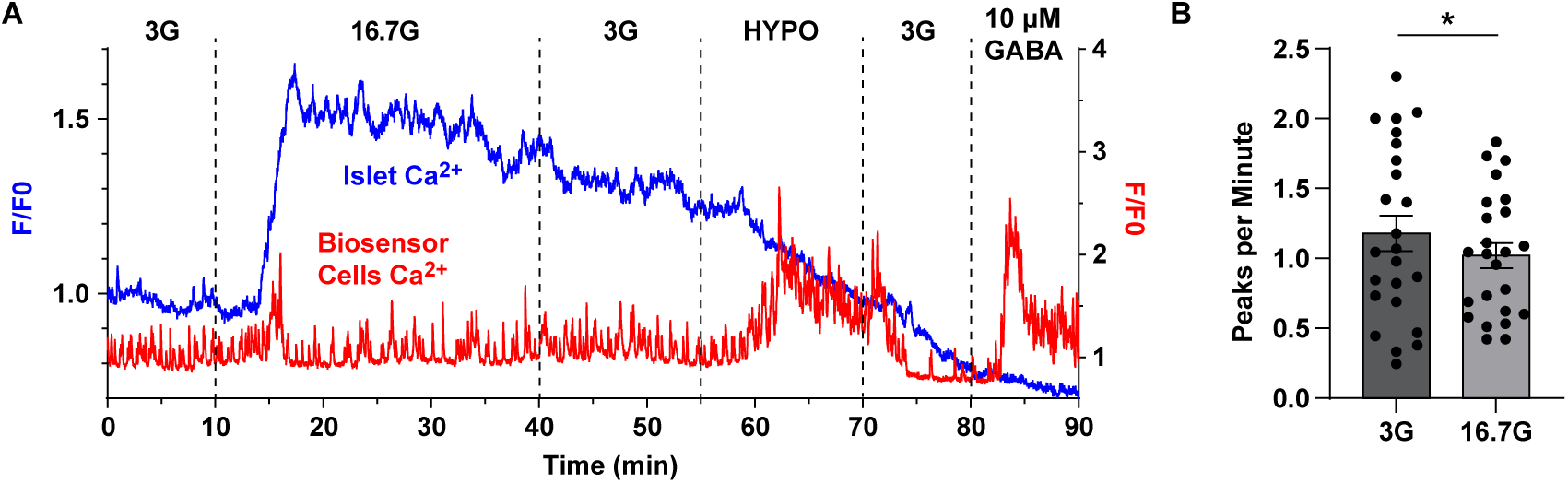
GABA Secretion Dynamics from Human Islets Differs from Mouse Islets. (**A**) Representative trace of islet Ca^2+^ and average biosensor cell Ca^2+^ for islet from one human donor. **(B)** Peaks per minute in 3 mM glucose (3G) and 16.7 mM glucose (16.7G) (n = 24, 3 donors, ±SEM, paired t-test, * p < 0.05).

After seeing this alignment between GABA release and islets [Ca^2+^]_i_ oscillations, we investigated how manipulating the [Ca^2+^]_i_ response of the islet would affect the GABA secretion pulses. C57BL/6J mouse islets were treated with tolbutamide, diazoxide, or left untreated during high glucose stimulation (Figure 6D-F). The control islets have an initial [Ca^2+^]_i_ increase and then oscillate in high glucose (Figure 5D). The diazoxide-treated islets have no response to glucose and do not generate any [Ca^2+^]_i_ oscillations (Figure 6E) while the tolbutamide-treated islets have a sharp initial [Ca^2+^]_i_ increase that remains at a constant high [Ca^2+^]_i_ level with no oscillations (Figure 6F). The high glucose stimulation was split into first and second phases for analysis based on the initial [Ca^2+^]_i_ rise of the islet and then the remaining plateau and/or oscillations (Figure 6G-J). In the diazoxide-treated islets, no [Ca^2+^]_i_ rises are present in either phase, and the GABA release events with the biosensor cells were lost in both the rise and plateau phases (Figure 6G-H). In the tolbutamide-treated islets, no difference was observed in islet [Ca^2+^]_i_ or GABA release events in the rise phase, as the islets had a [Ca^2+^]_i_ increase in both conditions (Figure 6I-J). When the tolbutamide blocked the islet oscillations during the plateau Ca^2+^ phase, the GABA release events decreased significantly compared to the control (Figure 6J). These results demonstrate that when islet [Ca^2+^]_i_ is actively changing up or down, the GABA biosensors record GABA release, but when the islet [Ca^2+^]_i_ is constantly high or low, the GABA biosensors record significantly fewer GABA release events. Additionally, the metabolism of glucose alone does not seem to trigger the GABA release when islet electrical activity is blocked with diazoxide. This suggests a mechanism where the LRRC8A/D mediated GABA release pulses depend on cyclical beta cell depolarization events.

### GABA secretion dynamics from human islets differs from mouse islets

We used our biosensor cell array to investigate if GABA secretion dynamics in human islets (Figure 7A-B, Sup. Fig. 8). Human islets showed periodic GABA secretion events in both low and high glucose. Human islets exhibited a smaller, less-pronounced, first-phase glucose-induced GABA secretion event and a large GABA bolus following hypotonic stimulation. In contrast to mouse islets, the frequency of human islet GABA secretion pulses was relatively glucose insensitive and even decreased slightly, consistent with previous results^25^. Approximately 50% of recorded human islets did not exhibit a first-phase glucose-induced GABA release. These observations were consistent across three different donors. When quantified, the number of GABA pulses per minute was slightly but significantly reduced by about 10%, whereas in mouse islets the number of pulses per minute significantly increased during glucose stimulation. Measurement of the alignment of GABA pulses with Ca^2+^ oscillations was not performed in human islets because they lack the prominent Ca^2+^ peaks that are readily observed in mouse islets. Thus, the human islet GABA pulse pattern differs from mouse islets.

**Figure 8:**
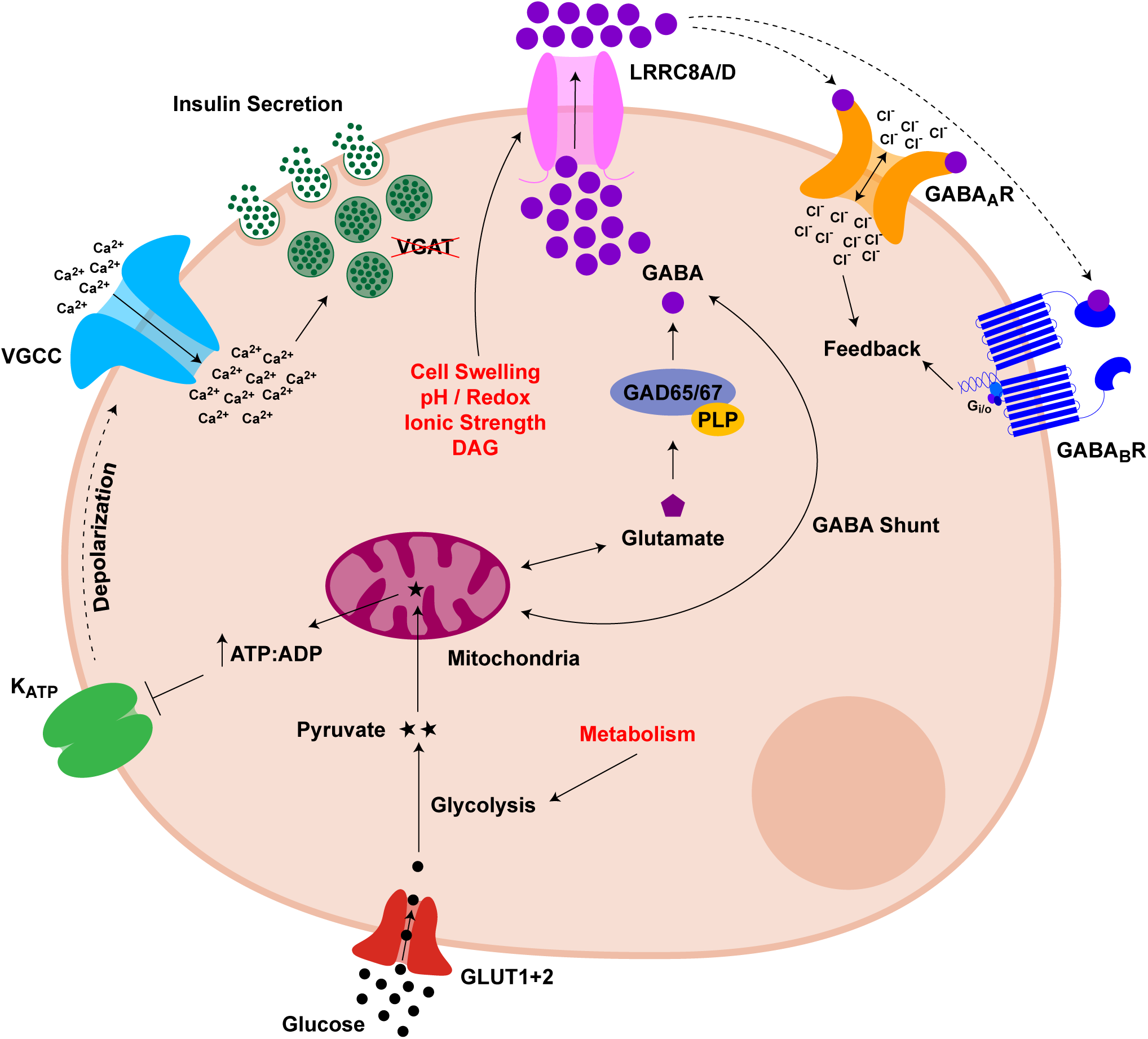
Proposed Mechanism for GABA Secretion from Beta Cells. Glucose is imported into the beta cell via GLUT1 and 2 glucose transporters. Glucose enters glycolysis and is transformed into two pyruvates. Pyruvate is transported into the mitochondria where it enters the citric acid cycle and produces ATP. The increase in intracellular ATP to ADP ratio inhibts the K_ATP_ channel. Closing this channel depolarizes the beta cell, opening the voltage-gated calcium channel (VGCC) which stimulates insulin release (without VGAT on the granules). VRAC is activated by potentially one or multiple previously published mechanisms of cell swelling, changes in pH or reduction/oxidation, changes in ionic strength, diacylglycerol (DAG), and beta cell metabolism. This VRAC activation of the LRRC8A/D VRAC channel releases GABA. The intracellular GABA is produced from glutamate with the glutamate decarboxylase 65 and/or 67 enzyme with co-factor pyridoxal-phosphate (PLP). GABA can also be metabolized back into the citric acid cycle via the GABA shunt with the enzyme GABA transaminase. The secreted GABA will then act on both the GABA_A_ receptor and GABA_B_ receptor, providing feedback to the beta cell through a chloride current and G-protein respectively.

## Discussion

By combining high-temporal-resolution biosensing with biochemical measurements, this study shows that GABA secretion from pancreatic islets is coupled to beta cell electrical activity but is separate from insulin secretion. A central finding is that GABA release occurs as discrete pulses that are in-phase with intracellular Ca²⁺ oscillations, indicating that islet rhythmic activity gates GABA release. This study also defines the LRRC8A/D isoform of VRAC as a principal pathway for GABA release. We do not interpret our data to suggest that Ca²⁺ directly activates VRAC. Instead, membrane depolarization and Ca²⁺ influx establish the cellular conditions permissive for VRAC activation and GABA efflux.

Our data are consistent with a model where GABA is released in discrete pulses that require oscillatory, rather than sustained, Ca²⁺ dynamics. We posit that these GABA pulses are superimposed over a steady, tonic secretion that is proportional to the rate of intracellular GABA biosynthesis governed by GAD enzyme activity. Tolbutamide-treated islets held in a continuously activated state without Ca^2+^ oscillations showed only a single GABA release event during the first-phase Ca^2+^ response. In the second phase, tolbutamide treated islets had few GABA release events, suggesting that GABA release pulses depend on Ca^2+^ oscillations rather than simply Ca^2+^ activation. The idea that discrete GABA release events are triggered during [Ca^2+^]_i_ transitions makes sense given that VRAC activation follows a rapid change in osmotic pressure or ionic strength. GABA pulses during Ca^2+^ waves also make intuitive sense if GABA acts to provide signaling feedback to the islet. The defective Ca^2+^ oscillations observed in GAD knockout mouse islets reinforces this hypothesis^64^. To further prove the idea that GABA pulses coincide with Ca^2+^ waves, we treated islets with diazoxide to hold open the K_ATP_ channels and prevent electrical depolarization and Ca^2+^ entry. All GABA release events, including the first-phase pulse, were eliminated. The diazoxide experiment shows that glucose entry and metabolism alone are not sufficient to trigger islet GABA pulses and suggests membrane depolarization is upstream of VRAC-mediated GABA release.

Diazoxide experiments also revealed an apparent discrepancy between measurements of GABA secretion at different temporal resolutions. While diazoxide completely abolished GABA release events detected by biosensor cells, it had no effect on the cumulative GABA accumulation measured by HPLC over longer, one-hour incubations. A tonic component of GABA release must persist in the absence of oscillatory activity that dominates the bulk measurements over longer timescales. This likely reflects a constitutive efflux governed by intracellular GAD enzyme activity and the size of the intracellular GABA pool. This tonic secretion, initially described by Braun et al.^6^, likely sits below the detection threshold of the biosensor cells, but adds up to substantial GABA accumulation over longer frames over which GABA pulses are superimposed. This distinction explains how GABA release can appear Ca²⁺-dependent in high-temporal-resolution measurements yet largely Ca²⁺-independent when assessed as cumulative output. Overall, these findings highlight that conclusions regarding islet GABA regulation are highly dependent on the temporal resolution of the measurement and helps to explain the lack of consistent observations in the literature.

Our results also address longstanding discrepancies regarding glucose regulation of GABA secretion. Earlier work, including our own, concluded that GABA release is largely glucose-independent while others have shown the GABA release is triggered by glucose and is coupled to Ca^2+^ entry^6, 24, 25^. Cumulative islet GABA release not being correlated with glucose stimulation is consistent with biochemical measurements by Baekkeskov, Caicedo and Phelps^25^, Pipeleers^67, 68, 69^, Tamarit-Rodriguez^70, 71, 72^, Roper ^73^, and Naji^74^. Cumulative GABA release appears to track more closely with overall intracellular GABA levels rather than glucose concentration. These discrepancies can now be understood to depend on both the species of the islet and the timeframe of the measurement. By resolving secretion dynamics at low– and high-temporal-resolution, and short and long timeframes, we identify a rapid, glucose-triggered GABA pulse that occurs immediately following the transition to high glucose and coincides with the first-phase Ca²⁺ response. This acute glucose-induced GABA release in rodent islets was confirmed by four different observations: biosensor cell measurements, iGABASnFR, HPLC of short static incubations, and analysis of perifusion fractions. This acute release is also consistent with the previous observations by Braun et al., showing glucose and Ca^2+^ entry gate islet GABA release but now we show that this GABA release is short-lived. This acute GABA secretion component is followed by additional oscillation-coupled pulses during the second phase. In contrast, measurements of cumulative GABA secretion over extended periods show no positive glucose dependence and instead reflects a gradual depletion of intracellular GABA pools during high cellular respiration and/or a reduction in GAD enzyme activity. These findings demonstrate that GABA secretion is glucose-dependent on short timescales but becomes glucose independent at steady state. The observed decrease in intracellular GABA following prolonged glucose stimulation is consistent with previous reports of increased utilization of GABA in the tricarboxylic acid cycle via the GABA shunt pathway^70^. In brain tissue extracts, GAD enzyme activity has been shown to be inhibited by ATP and protein phosphorylation (e.g. PKA) and increased by Ca^2+^-dependent dephosphorylation (e.g. calcineurin)^75, 76^. It would thus be informative to investigate the short– and long-term dynamics of GAD enzyme activity in beta cells during transition from low to high glucose in future studies.

In our study, human islets showed no positive glucose-dependence on the frequency of GABA pulses or cumulative GABA secretion, in agreement with our previous observations in human islets^25^. A small, acute glucose-induced GABA release was observed in some human islet biosensor cell traces but was absent from other traces, suggesting the glucose-induced GABA release is more subtle in human islets than in mouse islets. One explanation for this could be due to species differences in the isoform of GAD enzyme expressed, where mouse beta cells express GAD67 and human beta cells express GAD65. GAD enzyme activity is regulated by a cycle of activation and inactivation determined by binding of the co-factor, pyridoxal 5’-phosphate (PLP). Most GAD65 largely exists as an inactive apoenzyme and GAD67 is mainly active holoenzyme due to strong kinetic differences in the deactivation rate between GAD isoforms (GAD65 auto-deactivates by separating from PLP fifteen times more rapidly than GAD67). Mouse islets may therefore have a larger pool of active GAD holoenzyme available to rapidly respond to glucose-driven increases in glutamate substrate, as glucose metabolism raises cytosolic glutamate through anaplerotic TCA flux and subsequent transamination of α-ketoglutarate.

We next asked if our observations of GABA release represent vesicular or tonic secretion mechanisms. Our data argues against a dominant role for vesicular co-release of GABA with insulin. Although previous studies have suggested that a subset of insulin granules may contain GABA, we find that large bidirectional changes in insulin secretion induced by forskolin or diazoxide do not produce corresponding changes in net GABA release. These findings indicate that GABA secretion is not tightly coupled to insulin exocytosis and is unlikely to arise primarily from insulin-containing large dense-core vesicles. Consistent with a non-vesicular mechanism, we find no evidence for expression of the vesicular GABA transporter VGAT in beta cells using reporter mice and transcriptomic and proteomic datasets. These observations resolve conflicting reports in the literature and argue against classical synaptic-like vesicular release. Although VMAT2 has been proposed as a low-affinity alternative transporter and is expressed in beta cells, recent findings argue against its ability to transport GABA under physiological conditions^30, 31, 32, 33^.

Our results support a model in which GABA is released directly from the cytosol through the LRRC8A/D isoform of VRAC. Genetic disruption of LRRC8D, and previously LRRC8A, markedly reduced GABA release pulses and hypotonic-induced GABA secretion, confirming the involvement of this channel. Because hypotonic stimulation is unlikely to occur under physiological conditions, we should consider potential physiological triggers of VRAC. Prior work demonstrated that LRRC8A-dependent Cl^-^currents in beta cells are activated by glucose metabolism driven cell swelling resulting from intracellular accumulation of glucose metabolites. Our results with diazoxide suggest that glucose metabolism alone, in the absence of membrane electrical activity, is insufficient for LRRC8A/D GABA secretion, but could drive Cl^-^ efflux through the other VRAC isoforms. Known modulators of VRAC include changes in cell volume, redox state, and ionic strength. One hypothesis for physiological LRRC8A/D gating that we find compelling is changes in intracellular Cl^-^. Beta cells maintain a high intracellular Cl^-^ due to high NKCC1 expression. A proposed mechanism linking glucose-stimulated Ca²⁺ signaling to non-vesicular GABA release involves Ca²⁺-activated Cl⁻ channels (CaCCs), particularly ANO1, which is functionally expressed in pancreatic beta cells^77^, or BEST1^78^, which is expressed in islets according to public proteomics and RNA sequencing datasets. During glucose stimulation, oscillatory Ca²⁺ influx through voltage-gated Ca²⁺ channels could directly activate ANO1 or BEST1, driving Cl⁻ efflux and transiently reducing intracellular [Cl⁻]. In addition, other LRCC8A VRAC heteromers and cAMP-sensitive CFTR channels might also contribute to glucose-induced Cl^-^ efflux. Given that VRAC channels formed by LRRC8A/D heteromers are sensitive to reduced intracellular ionic strength^39^, each Ca²⁺ oscillation could generate a corresponding cycle of Cl⁻ efflux that de-inhibits VRAC, permitting a pulse of GABA release in phase with the Ca²⁺ oscillation. This model would explain the observed synchrony between glucose-induced Ca²⁺ transients and GABA secretion. However, the extent to which Cl⁻ efflux can overcome NKCC1-mediated Cl⁻ re-accumulation to produce a net reduction in intracellular Cl⁻ sufficient to gate VRAC remains to be directly demonstrated, and future experiments using real-time Cl⁻ imaging alongside GABA release measurements and genetic and pharmacological models will be necessary to test this hypothesis.

We suggest a model where VRAC activation is coordinated with beta cell depolarization and Ca^2+^ mobilization into the beta cell cytosol, creating conditions that are permissive for GABA release via the LRRC8A/D channel (Figure 8). Known VRAC activators such as cell swelling in high glucose^40, 42, 79, 80, 81, 82^, changes in cytosolic ionic strength^39^, changing pH and oxidation-reduction (redox) potential^83^, and diacylglycerol (DAG) protein kinase C (PKC) activity at the plasma membrane^84^ could each also contribute. Once a pulse of GABA is secreted during a Ca^2+^ peak, GABA provides feedback through GABA_A_ and GABA_B_ receptors in endocrine cells to help synchronize the islet and prepare it for the next oscillation. In this manner, GABA may act as a diffusible autocrine and paracrine signal that reinforces the islet oscillatory rhythm.

We wish to acknowledge several limitations of this study. While the GABA biosensor cell system provides high temporal resolution and sensitivity, it is an indirect readout of extracellular GABA and is prone to influence by local flow, convection, and diffusion gradients, biosensor cell positioning relative to the islet, and biosensor internal cellular activities and variability in receptor expression. The biosensor cell system works best for detecting rapid changes to ambient GABA and may be inadequate for measuring steady-state, tonic GABA secretion given that it is most sensitive between 1 µM and 10 µM, while steady-state interstitial concentrations of GABA in the islet may be closer too 100 nM^20^. Furthermore, species differences between mouse and human islets, including differences in Ca²⁺ dynamics and GAD isoform expression, limit the generalizability of our findings in mouse islets to human physiology. An additional species limitation is that we used different glucose stimulating conditions for mouse and human islets. The reason for this difference is that we found mouse islets tended to generate the best and most predictable Ca^2+^ oscillations when using 5 and 11 mM glucose. Changing the mouse islet glucose bounds higher or lower resulted in poorer, less predictable oscillations. In contrast, human islets generated the best dichotomous Ca^2+^ response using 3 and 16.7 mM glucose. Human islets in our hands also generally lacked the deep slow-wave Ca^2+^ oscillations characteristic of C57BL/6J islets, which is not unusual for human donor islets. While we propose a model linking Ca²⁺-oscillations to VRAC gating, clearly GABA is still released in the absence of oscillations. Thus, oscillations may coordinate timing VRAC gating but are not a hard requirement for all GABA release. Finally, while VRAC acts fundamentally as an ionic strength sensor, ionic strength depends on all ions. Intracellular ion dynamics such as Cl⁻ were not directly measured and remain to be experimentally validated. Changes in ionic strength and/or cell swelling that gate VRAC under physiological conditions remain a topic of active investigation.

## Author Contributions

A.E.S. planned and performed the experiments, analyzed the data, wrote Matlab code for analysis, and wrote the manuscript. C.S.L. helped perform Ca^2+^ imaging with islets and biosensor cells and helped isolate islets. S.M.F. provided *Gad* ^βKO^ and Ins-cre mice and isolated their islets. A.E.C. performed single cell RNA sequencing and protein expression analysis in R. D.S. helped perform biosensor cell experiments. D.W.H. performed western blot of *Gad* ^βKO^ and Ins-cre islets and HPLC analysis of GABA content in *Gad* ^βKO^ and het islets. T.N. and S.P.G. provided pancreas and brain tissues from Vgat-ires-cre x Ai14 tdTomato mouse. E.S. and T.P.S. performed Ca^2+^ FLIPR analysis for GABA biosensor cells and funded that portion of the study. E.A.P. conceived and designed the study, acquired funding, supervised the research, and edited the manuscript.

## Data and Code availability

Matlab codes used for Ca^2+^ analysis are available at: https://github.com/PhelpsLabUF/Stis-et-al.-2025/ Authors will provide raw data upon request.

## Ethical approval

Human pancreatic islets were obtained from deceased non-diabetic donors from the NIDDK-funded Integrated Islet Distribution Program (IIDP) at City of Hope and from Prodo Laboratories, Inc. Cadaveric human islets for research were approved as non-human by the University of Florida Institutional Review Board (IRB No. NH00041892). All experimental protocols using mouse islets were approved by the University of Florida Institute Animal Care and Use Committees (IACUC No. 202400000147).

## Acknowledgments

We wish to thank Louis Scampavia, Senior Scientific Director, Department of Molecular Medicine, The Herbert Wertheim UF Scripps Institute for Biomedical Innovation & Technology for assistance with FLIPR assays. We also wish to thank Andrece Powell and Alma Peña Briseño for their assistance with human islet culture and lab logistics.

Human pancreatic islets and/or other resources were provided by the NIDDK-funded Integrated Islet Distribution Program (IIDP) (RRID:SCR_014387) at City of Hope, NIH Grant # 2UC4DK098085 and the JDRF-funded IIDP Islet Award Initiative.

This work was funded by the following NIH grants: T32DK108736 (A.E.S.), P01AI042288 (E.A.P.), R01DK132387 (E.A.P.), R01DK124267 (E.A.P.), R01DK123292 (E.A.P.), the NIDDK-supported Human Islet Research Network (HIRN, RRID:SCR_014393; https://hirnetwork.org) UH3DK122638 (E.A.P.), Breakthrough T1D 2-SRA-2023-1313-S-B (E.A.P.), Leo, Claire and Robert Adenbaum Foundation (E.A.P.), and Herbert Wertheim UF Scripps Institute for Biomedical Innovation & Technology Shared Services HTS Core (T.P.S.).

---

**Supplementary Figure 1:**
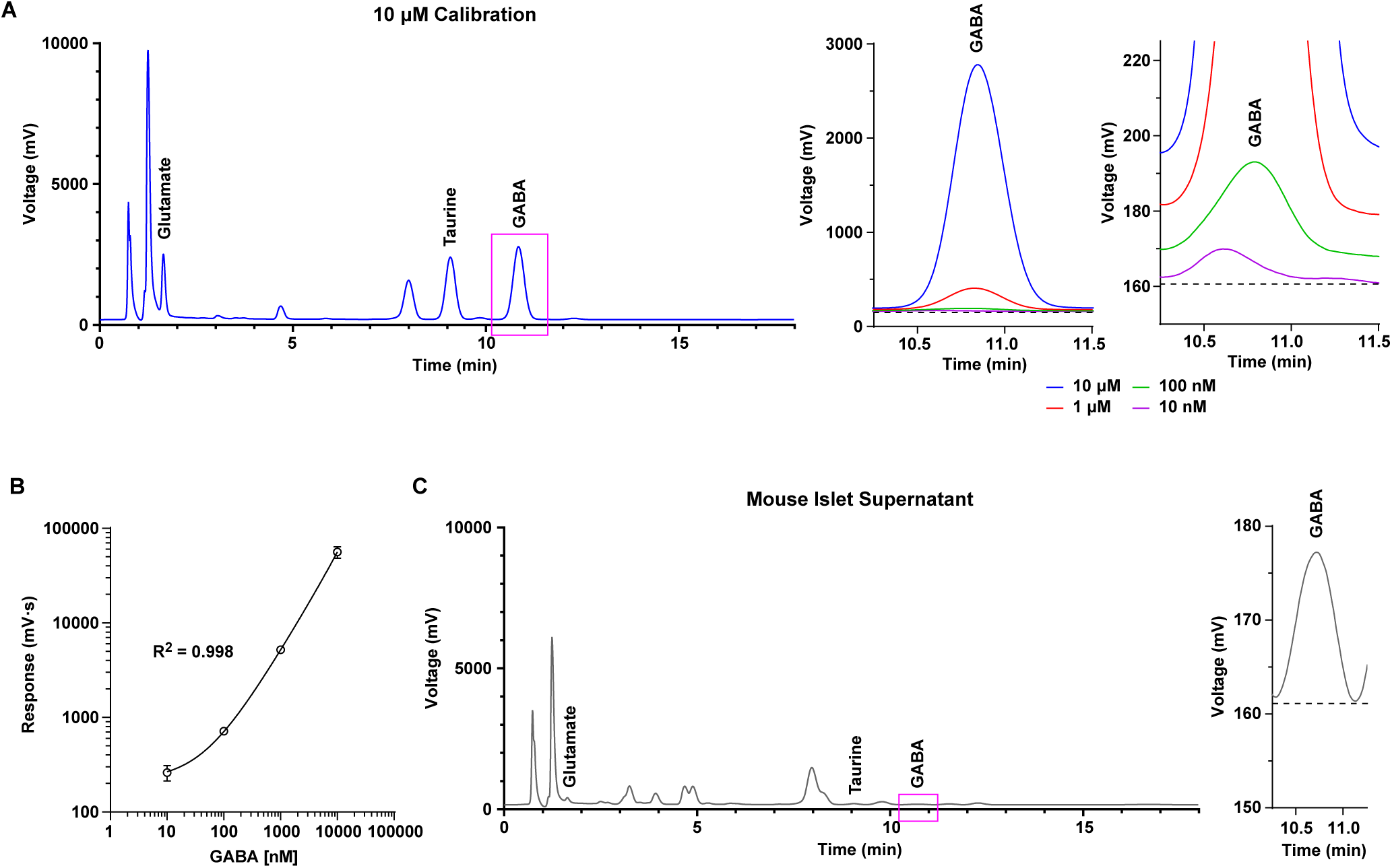
GABA Is Detected by HPLC. **(A)** (Left) HPLC trace of 10 μM glutamate, taurine, and GABA in KRBH buffer used for calibration. (Middle) Zoomed in trace of GABA peak for the 10 μM, 1 μM, 100 nM, and 10 nM GABA titration. (Right) More zoomed in trace to see 100 nM and 10 nM GABA titrations. **(B)** Calibration quadratic curve fit for GABA (95% CI, n = 3). **(C)** Sample trace of mouse islet supernatant from static incubation in 5G KRBH (Left). Zoomed in GABA peak from the trace (Right).

**Supplementary Figure 2:**
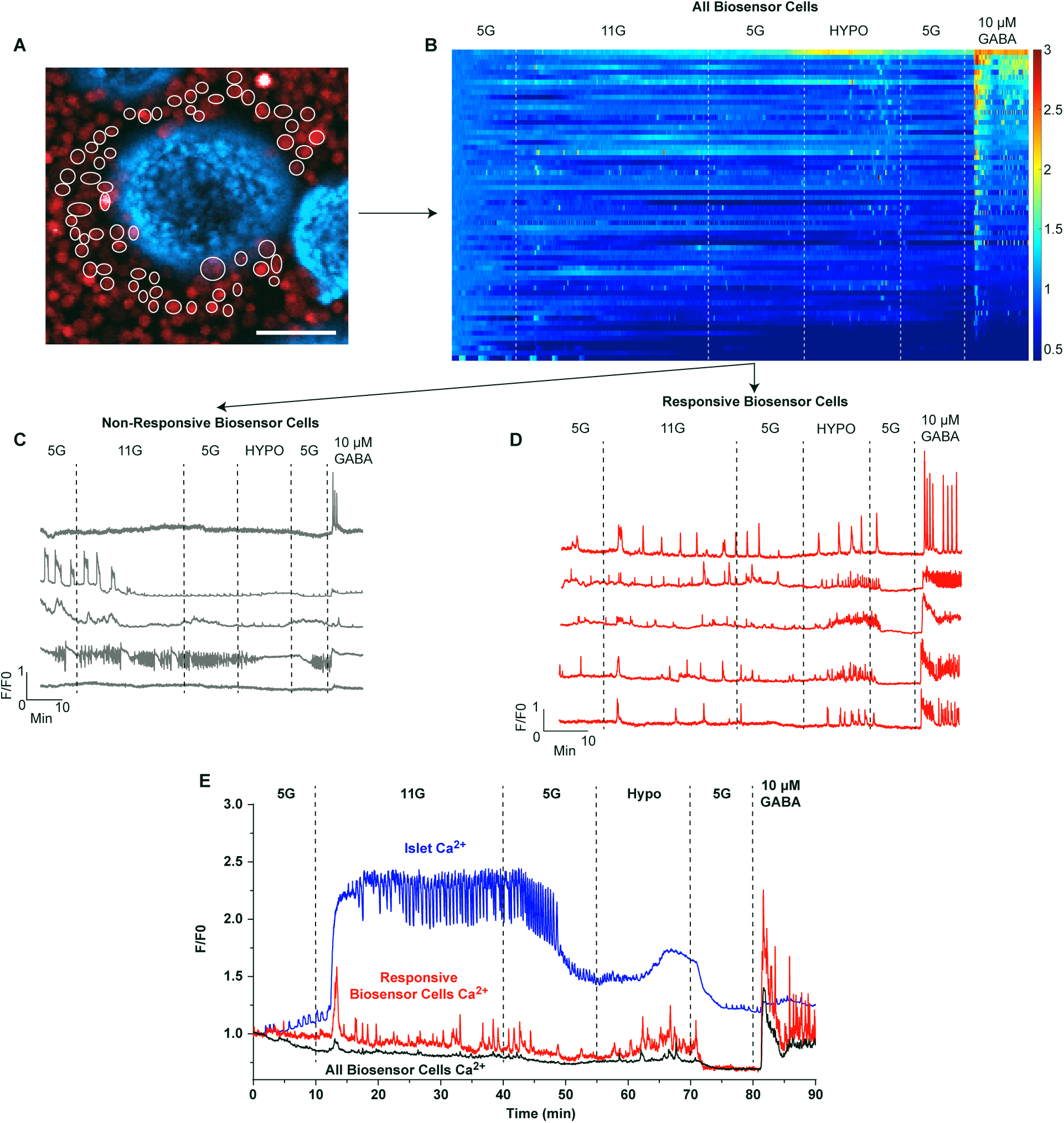
Analysis Pipeline of GABA Biosensor Cells from Ca^2+^ Recordings for an Individual Islet. **(A)** Zoomed in image of single islet and surrounding biosensor cells. Cells circled are are analyzed in (B). (Scale Bar = 100 μm). **(B)** Heatmap of all biosensor cell Ca^2+^ responses. **(C)** Example traces of non-responsive biosensor cells Ca^2+^ responses. **(D)** Example traces of responsive biosensor cell Ca^2+^ responses. **(E)** Trace of islet Ca^2+^, average responsive biosensor cell Ca^2+^, and average of all biosensor cell Ca^2+^.

**Supplementary Figure 3:**
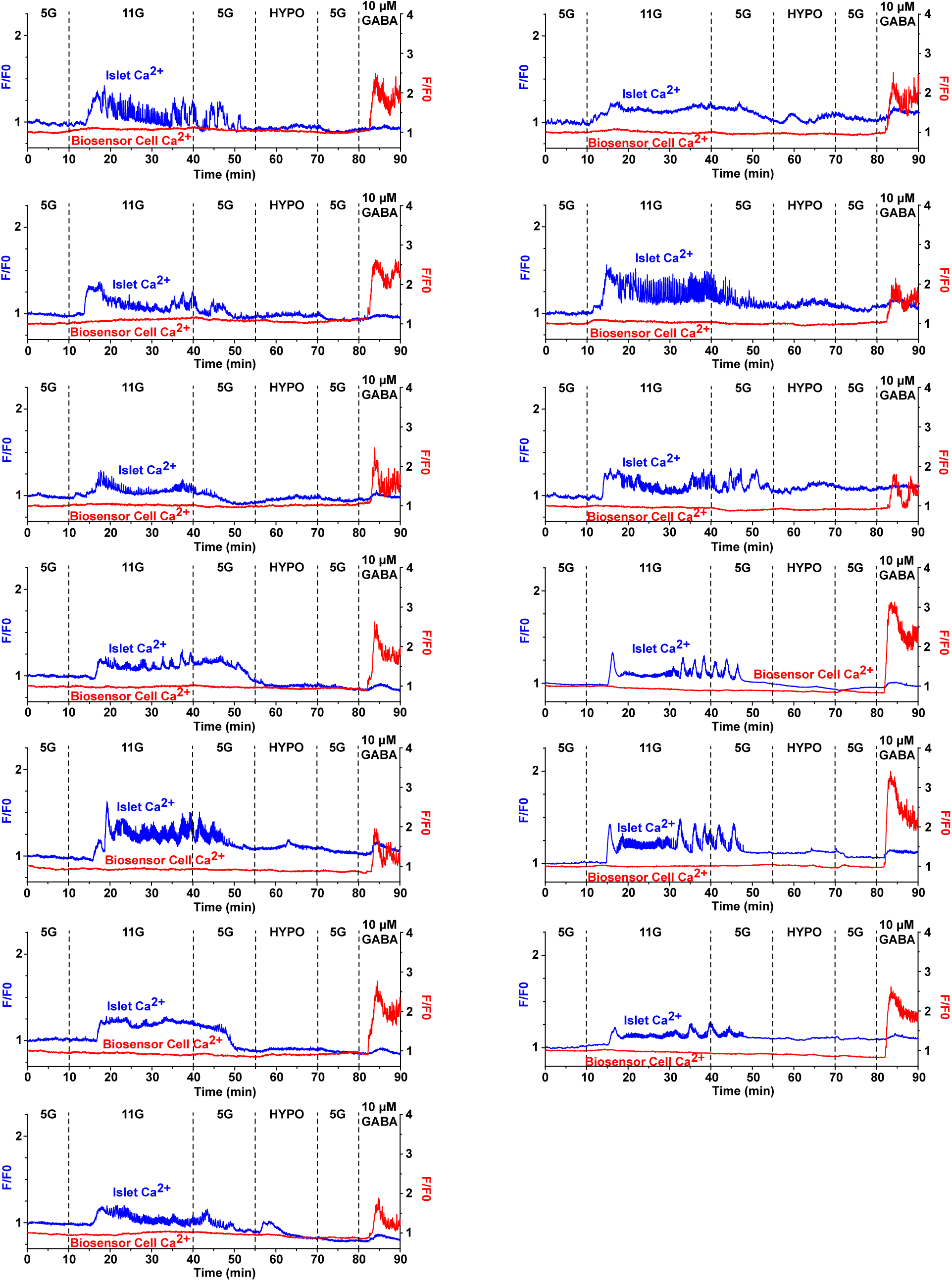
*Gad* ^βKO^ Traces. Additional *Gad* ^βKO^ Ca^2+^ traces from n = 3 mice on top of GABA biosensor cells Ca^2+^ traces.

**Supplementary Figure 4:**
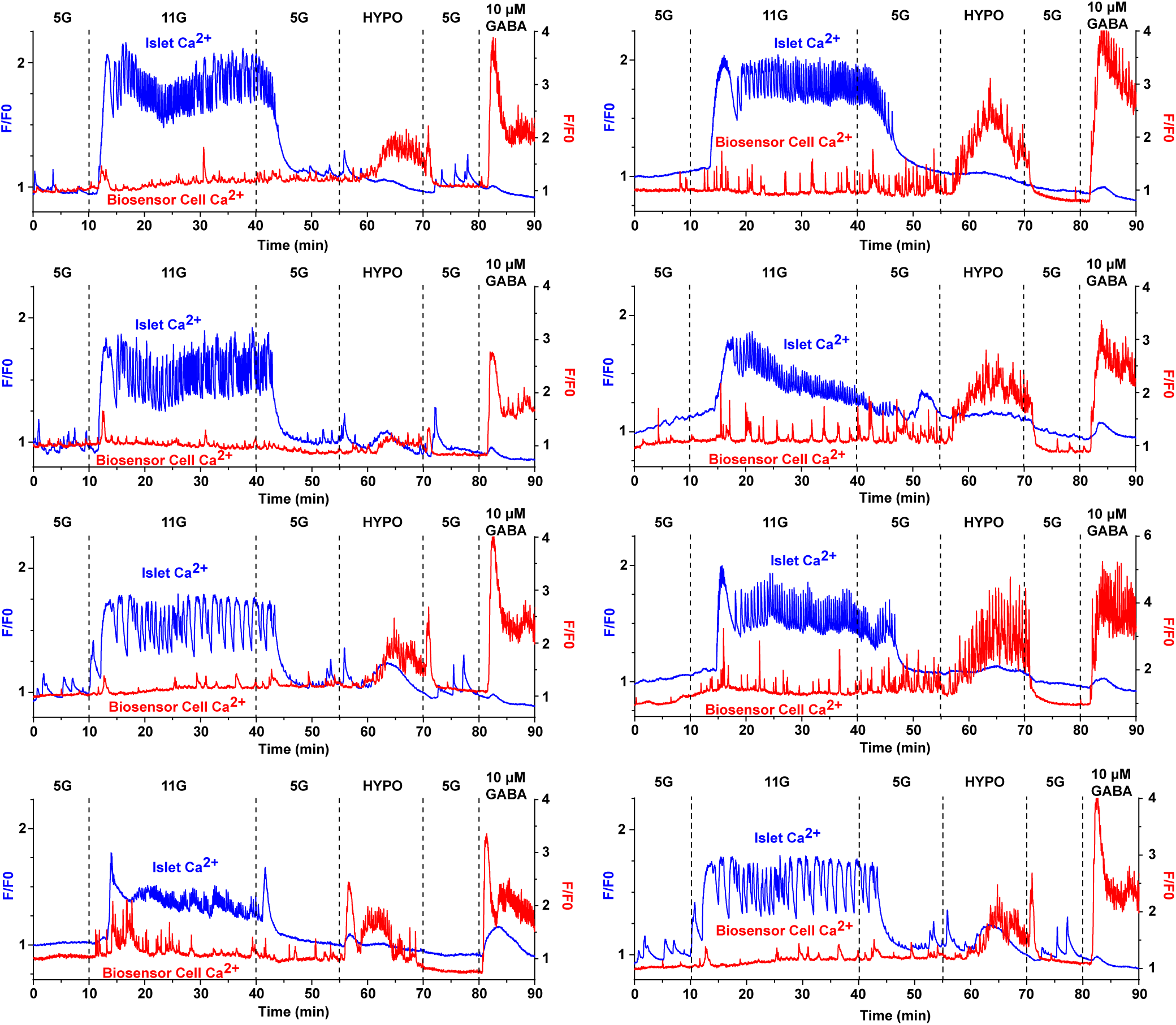
Ins-Cre Traces. Additional Ins-cre Ca^2+^ traces from n = 3 mice on top of GABA biosensor cells Ca^2+^ traces.

**Supplementary Figure 5:**
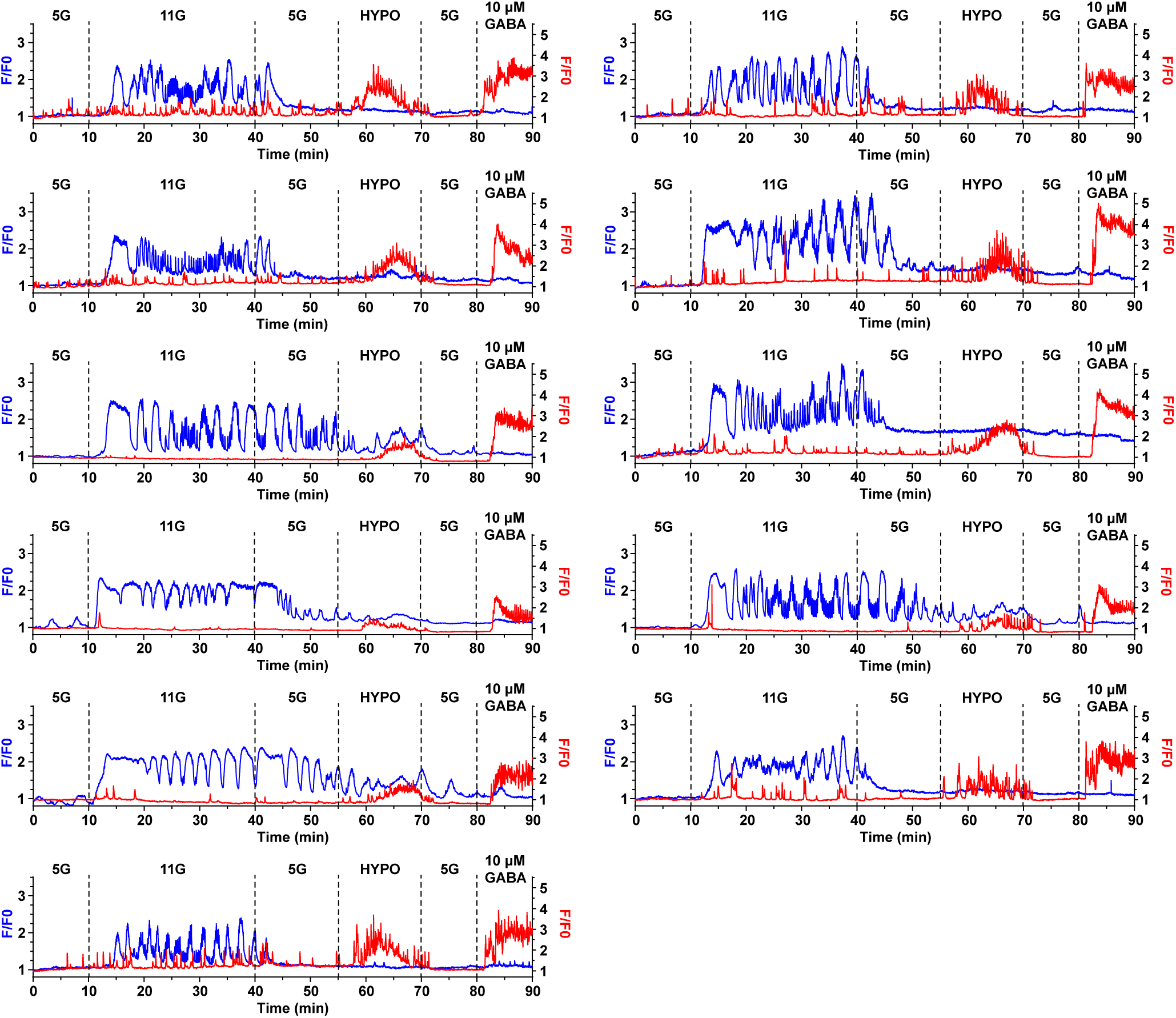
Scramble shRNA C57BL/6J Traces. Additional scramble shRNA C57BL/6J Ca^2+^ traces from islets (blue) of n = 3 mice on top of GABA biosensor cells Ca^2+^ traces (red).

**Supplementary Figure 6:**
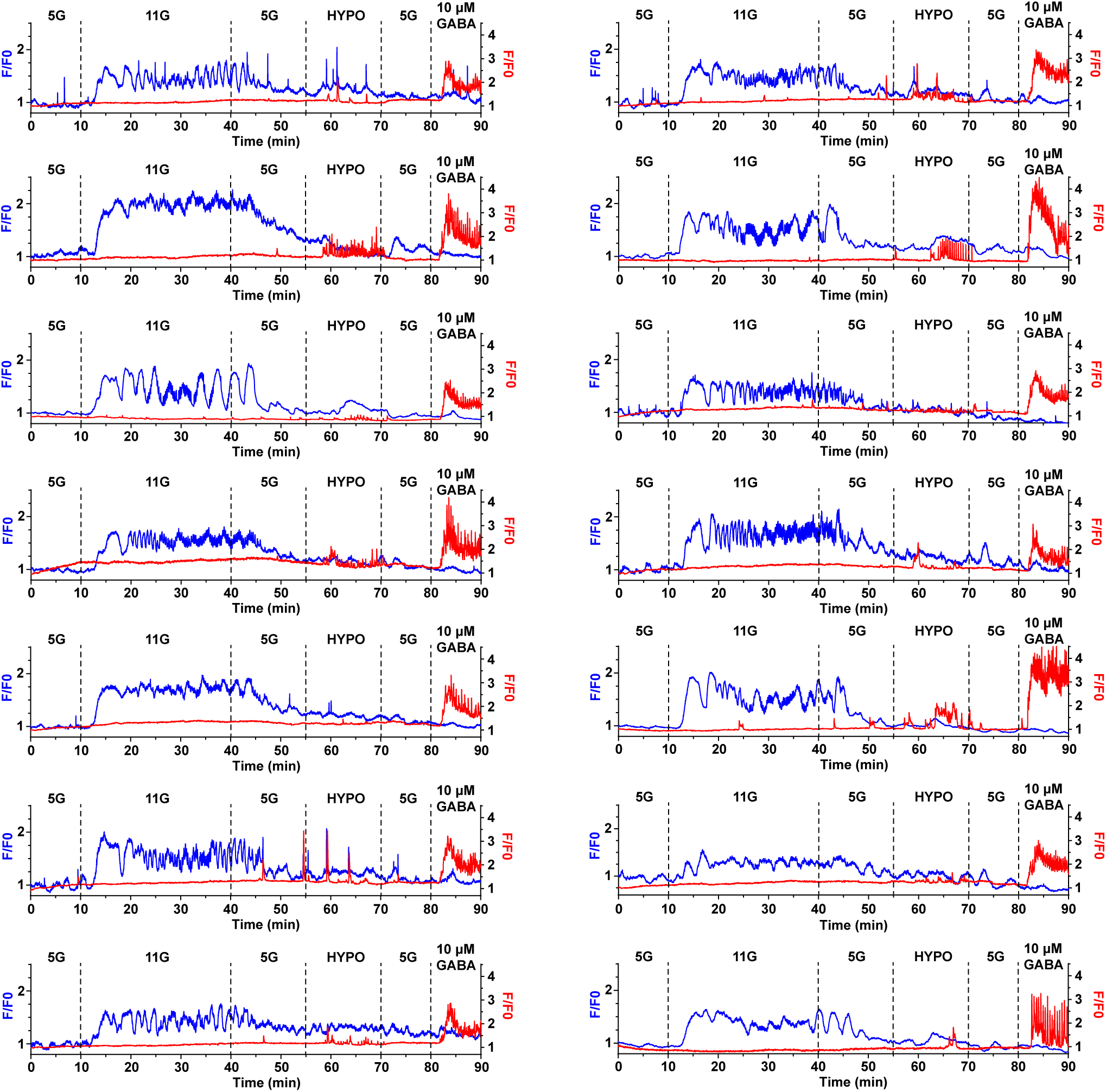
LRRC8D shRNA C57BL/6J Traces. Additional LRRC8D shRNA C57BL/6J Ca^2+^ traces from islets (blue) of n = 3 mice on top of GABA biosensor cells Ca^2+^ traces (red).

**Supplementary Figure 7:**
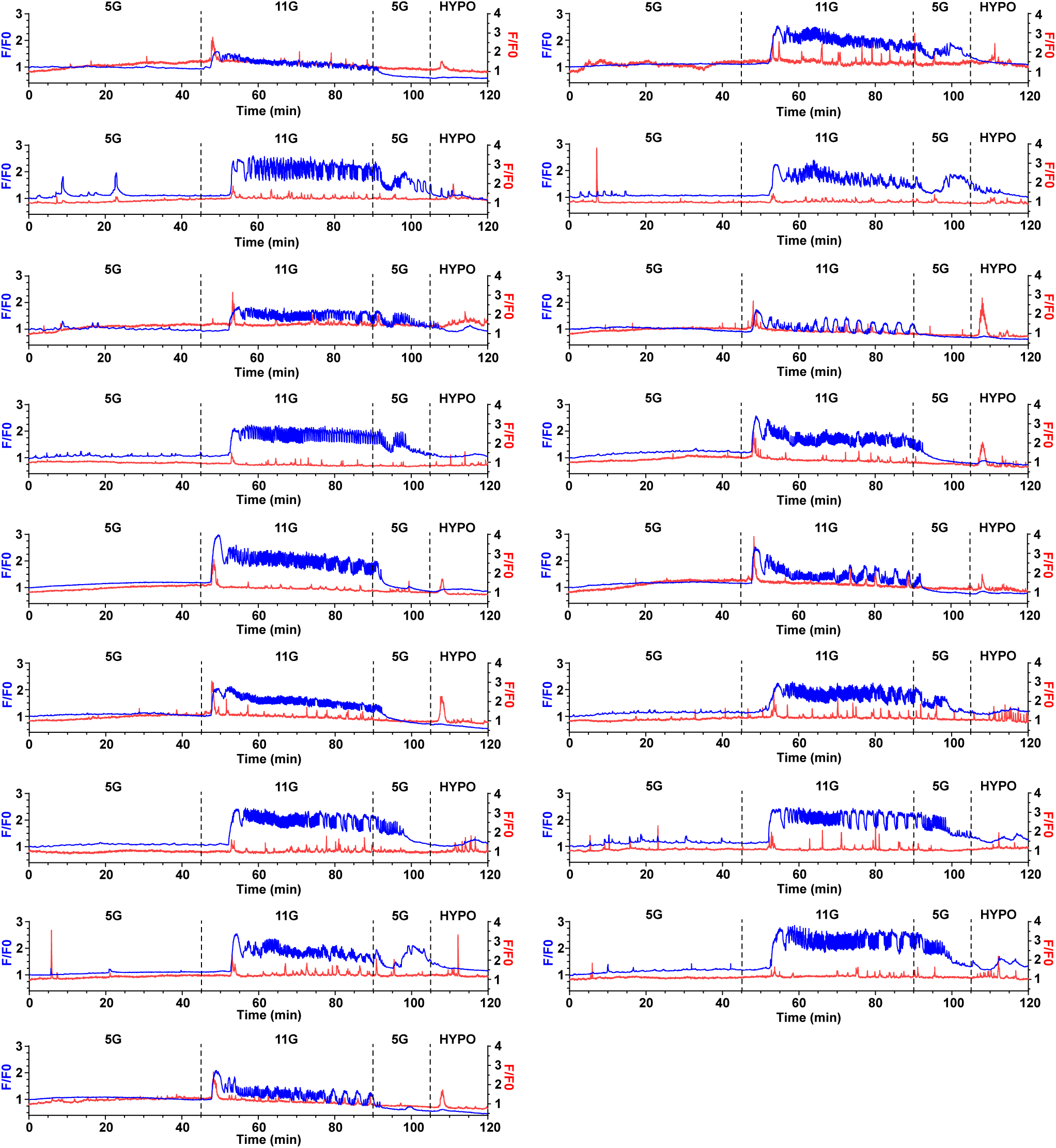
C57BL/6J Traces. Additional C57BL/6J Ca^2+^ traces from islets (blue) of n = 2 pooled mice on top of GABA biosensor cells Ca^2+^ traces (red).

**Supplementary Figure 8:**
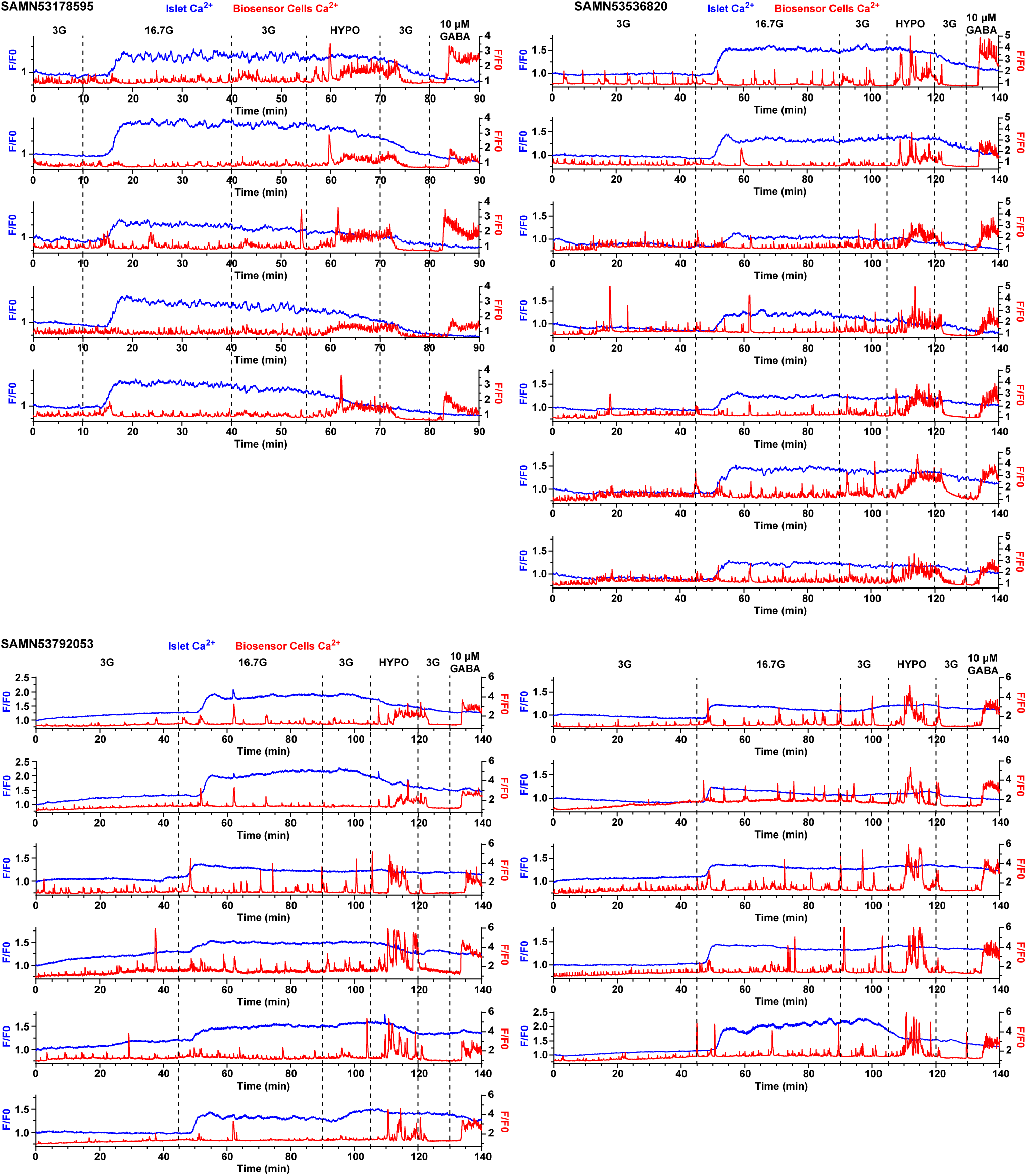
Additional Human Islet GABA Biosensor Cell Traces.

